# *Acinetobacter baumannii* represses type VI secretion system through an Mn^2+^-dependent sRNA-mediated regulation

**DOI:** 10.1101/2022.07.30.502137

**Authors:** Somok Bhowmik, Avik Pathak, Kuldip Devnath, Abhiroop Sett, Shivam Pandey, Nishant Jyoti, Timsy Bhando, Jawed Akhter, Saurabh Chugh, Ramandeep Singh, Tarun Kumar Sharma, Ranjana Pathania

**Affiliations:** Department of Biosciences and Bioengineering, Indian Institute of Technology Roorkee, Uttarakhand-247667, India; Translational Health Science and Technology Institute, Faridabad, Haryana-121001, India; Centre of Excellence in Disaster Mitigation and Management, Indian Institute of Technology Roorkee, Uttarakhand-247667, India

**Keywords:** Mn^2+^, oxidative stress, post-transcriptional regulation, small RNA, T6SS

## Abstract

Type VI secretion system (T6SS) is utilized by many Gram negative bacteria for eliminating competing bacterial species and manipulating host cells. *Acinetobacter baumannii* ATCC 17978 utilizes T6SS at the expense of losing pAB3 plasmid to induce contact-dependent killing of competitor microbes, resulting in the loss of antibiotic resistance carried by pAB3. However, the regulatory network associated with T6SS in *A. baumannii* remains poorly understood. Here, we identified an Mn^2+^-dependent post-transcriptional regulation of T6SS mediated by a bonafide small RNA, AbsR28. *A. baumannii* utilizes MumT (Mn^2+^-uptake inner membrane transporter) for the uptake of extracellular Mn^2+^ during oxidative stress. We demonstrate that the abundance of intracellular Mn^2+^ enables complementary base-pairing of AbsR28-*tssM* mRNA (that translated to TssM, one of the vital inner membrane components of T6SS), inducing RNase E-mediated degradation of *tssM* mRNA and resulting in T6SS repression. Thus, AbsR28 mediates a crosstalk between MumT and T6SS in *A. baumannii*.

**IMPORTANCE:** Small RNAs (sRNAs) are identified as critical components within the bacterial regulatory networks involved in fine regulation of virulence-associated factors. The sRNA-mediated regulation of *Acinetobacter baumannii*’s T6SS was unchartered. Our findings reveal a novel underlying mechanism of an Mn^2+^-dependent sRNA-mediated regulation of T6SS in *A. baumannii*. We show that binding of Mn^2+^ to AbsR28 aids in the complementary base-pairing of AbsR28-*tssM* mRNA, resulting in RNase E-mediated processing of *tssM* and T6SS repression. The findings also shed light on *A. baumannii*’s preference for antibiotic resistance over contact-dependent killing during infection.

## INTRODUCTION

*Acinetobacter baumannii* is a Gram-negative pathogen and has gained importance due to its ability to cause wound and burn infections, sepsis, meningitis, urinary tract infections, bloodstream infections, and ventilator-associated pneumonia (1–5). Treating hospital-acquired *A. baumannii* infections is a major hassle, as most *A. baumannii* clinical strains are multidrug-resistant (MDR) (6). Due to this severe global impact on public health, it is essential to identify the molecular machinery adapted by *A. baumannii* to escape from host-mediated immune response and establish pathogenesis.

One of the vital host-mediated immune responses encountered by pathogens is the phagocytic cell-mediated free metal ion limitation, termed as “host-mediated nutritional immunity” and generation of reactive oxygen species (ROS) at the site of infection (7, 8). Mn^2+^ sequestration by neutrophils at the infection site reduces the activity of bacterial metalloenzymes such as superoxide dismutase (SOD) and catalase, which bacteria use to break down ROS (7, 9, 10). To acquire Mn^2+^ during oxidative stress and host-mediated Mn^2+^ restriction, *A. baumannii* utilizes a high-affinity Mn^2+^ acquisition system MumT (<u>m</u>anganese and <u>u</u>rea <u>m</u>etabolism transporter) (11, 12).

A synergy between the type VI secretion system (T6SS) and Mn^2+^-transporter has been observed in *Burkholderia thailandensis* (13). *B. thailandensis* secretes an effector protein TseM through T6SS that scavenges Mn^2+^ from the extracellular milieu and delivers Mn^2+^ through an outer membrane transporter (MnoT) to compensate for the scarcity of intracellular Mn^2+^ under oxidative stress (13). T6SS is one of the secretion systems frequently utilized by Gram-negative bacteria to promote contact-dependent killing of competitors by injecting harmful toxins into it (14–17). Although Mn^2+^-uptake systems have been reported as one of the major virulence factors in pathogenic bacteria (18, 19), crosstalk between Mn^2+^-transporter and T6SS remains unknown in *A. baumannii*.

The biogenesis of T6SS assembly is an enormous and energetically expensive process for bacteria (20). As a contact-dependent system with specific cellular targets (21), the expression of T6SS might be precisely regulated transcriptionally and post-transcriptionally (22, 21). Transcriptional repression of T6SS by *tetR*-like regulators carrying by pAB3 plasmid or its derivative is observed in *A. baumannii* (23), whereas post-transcriptional regulation of T6SS has not been explored yet. Small regulatory RNAs (sRNAs) are identified as critical components within the bacterial regulatory networks involved in post-transcriptional regulation of virulence-associated factors. The sRNAs are mostly 50-500 nucleotides long, function as global regulators of numerous bacterial physiological processes, and play a crucial role in regulating several virulence factors (24, 25). The sRNAs base pair with their cognate mRNA and regulate the translational activity and/or the stability of that particular mRNA often with the assistance of the RNA chaperone, Hfq (26). The sRNA-mediated regulation of T6SS in *A. baumannii* remains unexplored.

In this work, we show that the breakdown in Mn^2+^-uptake by deleting *mumT* increases T6SS expression during oxidative stress. Moreover, the elevated level of intracellular Mn^2+^ by overexpressing MumT reduces T6SS expression. These observations indicate the existence of an Mn^2+^-dependent T6SS regulation in *A. baumannii*. We explored the crosstalk between MumT and T6SS with the aim of understanding the underlying mechanism of this crosstalk. Intriguingly, we identify an sRNA, AbsR28 (*<u>A</u>. <u>b</u>aumannii* <u>s</u>mall <u>R</u>NA <u>28</u>), that mediates the crosstalk between MumT and T6SS. Collectively, we show that the uptake of Mn^2+^ by MumT is necessary for AbsR28-mediated post-transcriptional regulation of *tssM* mRNA, leading to T6SS repression in *A. baumannii*.

## RESULTS

### *A. baumannii* cells that switch to T6+ are unable to withstand oxidative stress

The T6SS in bacteria is often found to be essential for bacterial pathogenesis as it modulates the host immune response during infection (27–30). In contrast, certain T6SSs seem to function as antivirulence mechanisms, as evidenced by increased pathogenicity in T6SS deficient mutants relative to wild-type bacteria (31, 32). Such involvement of *A. baumannii*’s T6SS during infection remains enigmatic. Oxidative burst by phagocytic cells serves as a primary mechanism for microbial eradication in the host during infection. To assess whether *A. baumannii* cells expressing T6SS (T6+) would have an advantage in survival during phagocytic cell-mediated killing compared to *A. baumannii* cells with silent T6SS (T6-), we co-incubated phagocytic cells either with wild-type (WT) T6- or WT cells that switched to T6+ cells and checked their survival. The WT T6- and WT cells that switched to T6+ cells were isolated from *A. baumannii* ATCC 17978 based on the secretion of T6SS-associated protein Hcp (<u>h</u>emolysin-<u>c</u>oregulated <u>p</u>rotein, a positive marker for expression of T6SS) using several rounds of Hcp-ELISA followed by western blot to detect secreted Hcp. The cells that secreted Hcp were considered as T6+ (WT T6+ cells after losing pAB3 plasmid which carries *tetR*-repressors for T6SS are referred as WT T6+ throughout the study) and the cells that showed no detectable Hcp in the cell-free supernatant were considered as T6- (WT T6-cells carrying pAB3 plasmid which carries *tetR*-repressors for T6SS are referred as WT T6-throughout the study) (Fig S1A). To our surprise, WT T6-cells exhibited ∼60-70% survival, whereas WT T6+ cells exhibited only ∼20-30% survival against phagocytic cell-mediated killing compared to untreated controls (Fig 1A and S1B). This observation suggests that WT T6+ cells cannot withstand phagocytic cell-mediated killing. As neutrophils induce oxidative stress by producing deleterious superoxide radicals during infection, the WT T6- and WT T6+ cells were grown in an LB medium supplemented with methyl viologen (MV) to induce superoxide radicals (33–35), and bacterial growth was measured at the indicated time points. The strains displayed almost equal growth in LB, however, upon addition of MV, WT T6+ cells exhibited a remarkable growth defect (Fig 1B). This reinforced our previous observation that WT T6+ cells display enhanced sensitivity to oxidative stress compared to WT T6-cells. Interestingly, There are two mixed variants of *A. baumannii* ATCC 17978, one variant has a 44 kb island in the genome (referred as +AbaAL44), and the other one is devoid of that 44 kb island (referred as -AbaAL44) (36). It was demonstrated that the presence of *katX* in AbaAL44 island confers resistance to *A. baumannii* strain carrying AbaAL44 island against H_2_O_2_-mediated oxidative stress (36). It was surprising that WT T6+ strain is sensitive to oxidative stress despite possessing AbaAL44 island, while WT T6-strain exhibits more resistance to oxidative stress despite lacking the AbaAL44 island suggesting that the growth defect is not dependent on AbaAL44 status (Fig S1C). Intracellular ROS generation via Fenton chemistry is a hallmark of oxidative stress response (37, 38). Thus, we measured intracellular ROS level in WT T6- and WT T6+ cells by growing them in either LB broth or LB broth supplemented with MV by measuring the fluorescence of 2’,7’-dichlorofluorescein (DCF). A significant increase in ROS level was observed in WT T6+ cells compared to WT T6-cells treated with MV (Fig 1C), suggesting that WT T6+ cells are more sensitive to oxidative stress due to higher intracellular ROS accumulation.

**Figure 1.**
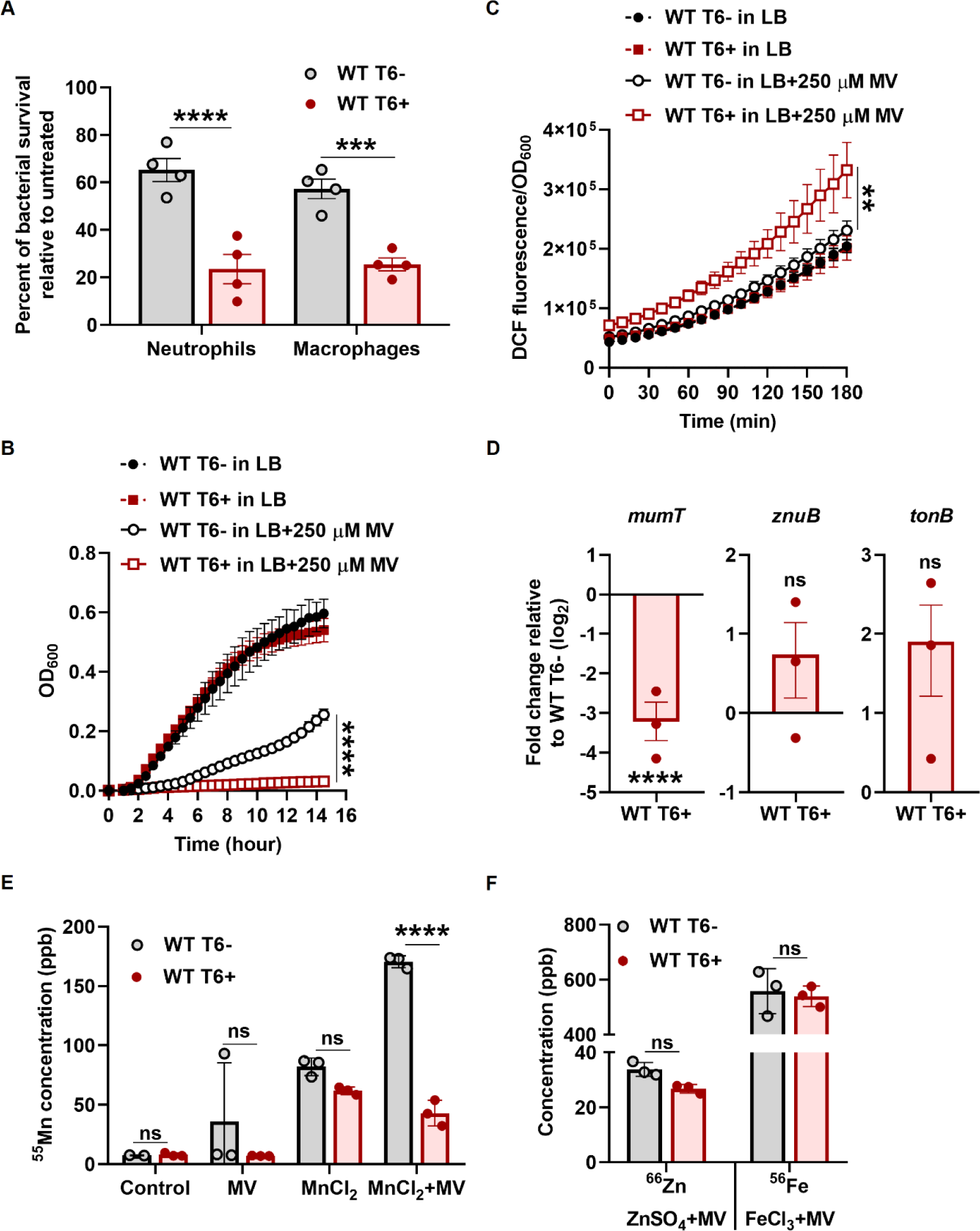
*A. baumannii* T6+ cells are defective in Mn^2+^-uptake and unable to cope with oxidative stress. (**A**) Bacterial cells were co-incubated with the phagocytic cells (human blood-derived neutrophils and macrophage RAW 264.7 cell line) for 4 h at an MOI of 1:1. The percentage of bacterial survival was enumerated by accounting for the respective untreated control (without phagocytic cells) as 100%. The data represent the mean of four independent experiments, each in biological triplicates ± SEM. (**B**) Growth of the indicated strains in the presence or absence of MV (250 µM) in LB broth. The data represents the mean of four biological replicates, each in technical triplicates ± SD. (**C**) Bacterial intracellular ROS generation was determined by measuring the fluorescence of DCF. The data represents the mean of four biological replicates ± SD. (**D**) Neutrophils were co-incubated with either WT T6- or WT T6+ cells, and the expression of *mumT*, *znuB*, and *tonB* in the bacterial cells was determined by qRT-PCR. The data represent the mean of three independent experiments, each in technical triplicates ± SEM. (**E and F**) Intracellular ^55^Mn, ^66^Zn, and ^56^Fe in cell pellet (OD_600_ normalized volume) were quantified by ICP-MS. The data represents the mean of three biological replicates ± SD. Statistical significance was determined using a multiple comparison two-way ANOVA test with the Sidak correction for multiple comparisons comparing the means of each group to one another (A, E, F) and Student’s t-test (B, C, D). ** denotes p-value <0.01, *** denotes p-value <0.001, **** denotes p-value <0.0001, ns denotes not significant.

### *A. baumannii* T6+ cells cannot withstand oxidative stress due to inadequate intracellular Mn^2+^

Bacteria utilize metal ion-dependent SODs (Mn/Fe SOD and Cu/Zn SOD) and catalases for breaking down intracellular ROS. Therefore, metal ion homeostasis governs the cellular ROS level. *A. baumannii* employs specialized metal uptake systems to import metal ions for the activity of these metalloenzymes during oxidative stress and to survive against host-mediated metal limitation (12, 39–41). Due to the compromised survival of WT T6+ cells in phagocytic cell-mediated killing, we hypothesized that WT T6+ cells may be defective in the uptake of free metal ions, which serve as cofactors for SODs and catalases required to break down ROS. To test this, we co-incubated neutrophils with either WT T6- or WT T6+ cells and checked the expression of *mumT*, *znuB*, and *tonB* genes involved in Mn^2+^-uptake, Zn^2+^-uptake, and Fe^2+/3+^-uptake, respectively, by quantitative reverse transcription PCR (qRT-PCR). There was no significant difference in the expression of *znuB* and *tonB* genes, but in the case of *mumT*, a ∼3.5-log_2_ fold reduction was observed in WT T6+ cells as compared to WT T6-cells (Fig 1D). Similarly, there was a significant fold reduction of *mumT* transcript levels in WT T6+ cells compared to WT T6-cells during oxidative stress when the bacterial cells were grown LB-medium supplemented with MV (Fig S1D). Due to the fold reduction of *mumT* expression in WT T6+ cells, we hypothesized that this could lead to intracellular Mn^2+^ deficiency in WT T6+ cells under oxidative stress. To examine this, we measured the concentration of intracellular Mn^2+^ using inductively coupled plasma mass spectrometry (ICP-MS). To determine the intracellular concentration of a specific metal ion, minimal media was preferred over complex media in this assay. The WT T6+ cells had ∼3-fold lower Mn^2+^ levels in cell pellets than WT T6-cells in the presence of MnCl_2_+MV (Fig 1E). In contrast, there were no significant differences in Zn^2+^ and Fe^2+/3+^levels in cell pellets of WT T6+ cells as compared to the WT T6-cells in the presence of ZnSO_4_+MV and FeCl_3_+MV, respectively (Fig 1F). Next, we hypothesized that the addition of sufficient MnCl_2_ might alleviate the oxidative stress toxicity in WT T6+ cells. The WT T6-strain grew almost equally in all conditions; supplementation of ZnSO_4_ and FeCl_3_ reverted the growth of WT T6+ cells to some extent, but supplementation of MnCl_2_ did not have much effect under oxidative stress (Fig S1E). Taken together, these data suggest that WT T6+ cells display impaired growth under oxidative stress due to inadequate intracellular Mn^2+^, which is crucial for *A. baumannii*’s SOD and catalase activity, and this defect in activity could not be restored even after supplementation of sufficient Mn^2+^ (Fig S1F, G). We isolated T6+ cells of two clinical strains of *Acinetobacter baumannii* (RPTC2 and RPTC3) and compared the growth of WT and T6+ cells of these isolates under oxidative stress. We observed that the T6+ strains of RPTC2 and RPTC3 showed a growth defect under oxidative stress compared to their respective WT strains (Fig S2A-C). Like *A. baumannii* ATCC 17978, RPTC2 and RPTC3 T6+ strains are more susceptible to neutrophil-mediated killing (Fig S2D).

### pAB3 plasmid has no role in mitigating oxidative stress

In view of the fact that WT T6-cells of *A. baumannii* ATCC 17978 switch to WT T6+ cells upon losing pAB3 plasmid (23) and the WT T6+ cells are sensitive to oxidative stress due to inadequate intracellular Mn^2+^, we speculated that the sensitivity might be due to the cells losing pAB3 plasmid which encodes several transcription factor regulators. To investigate this, we transformed pAB3 into the WT T6+ competent cells (which were devoid of pAB3), confirmed the transformants (Fig S3A, B), and assessed the growth of WT T6-, WT T6+, and WT T6+ cells transformed with pAB3 (referred as WT T6+/pAB3 throughout the study) strains in LB supplemented with MV. We observed that the presence of pAB3 does not confer resistance to oxidative stress as there is no growth difference between WT T6+ and WT T6+/pAB3 cells (Fig S3C). Next, we sought to determine whether *mumT* expression is reduced by T6SS functionality, rendering the T6+ cells defective in Mn^2+^ uptake (scenario 1), or if the opposite is true and a breakdown in Mn^2+^ uptake results in the T6+ phenotype (scenario 2) (Fig S3D). Consistent with the previous observation, we also observed that T6SS expression is repressed in WT T6+ strain upon transformation of pAB3 which contains *tetR*-repressors (Fig S3E) whereas the downregulation of *mumT* is unaffected in comparison to WT T6+ cells by the transformation of pAB3 into the WT T6+ cells (WT T6+/pAB3) (Fig S3F). This led us to infer that alteration of T6SS status does not impact *mumT* expression and ruling out the possibility of scenario 1.

### Breakdown in Mn^2+^-uptake results in an increase in T6SS expression

Mn^2+^ is an essential micronutrient for bacterial physiology, virulence, and survival against oxidative stress (18). *A. baumannii* utilizes an inner membrane transporter, MumT, for Mn^2+^ uptake during host-mediated metal restriction (12). To assess whether intracellular Mn^2+^ has any impact on T6SS regulation (scenario 2; Fig S3D), we first checked the effect of *mumT* on T6SS expression. To evaluate this, we created a deletion mutant of *mumT* in *A. baumannii* containing pAB3 plasmid (referred as Δ*mumT* throughout the study), grew both the WT T6- and Δ*mumT* cells under oxidative stress supplemented with MnCl_2_, and checked the expression of *hcp* by qRT-PCR. Hcp is a structural component of T6SS (Fig 2A). Interestingly, we observed a ∼6-log_2_ fold increase in *hcp* expression in Δ*mumT* cells with respect to WT T6-cells (Fig 2B). The data were further validated by Hcp-Western blot, where Hcp expression was highly induced in Δ*mumT* cells compared to WT T6-cells in both cell lysate (CL) and culture supernatant (S) (Fig 2C). We used a prey-predator assay to assess the T6SS-mediated killing of prey cells by WT T6- and Δ*mumT* cells, and the killing of prey cells confirmed the expression of T6SS. As expected, Δ*mumT* cells exhibited an efficient increase in the killing of prey cells compared to WT T6-cells (Fig 2D, S4A). We overexpressed *mumT* into WT T6+ and Δ*mumT* strains. We observed that the elevated intracellular Mn^2+^ (Fig S4B) due to an increase in MumT expression (Fig S4C) remarkably reduced Hcp secretion (Fig S4D) and T6SS activity compared to their respective control (Fig S4F). The percentage of Hcp-secretion positive cells increased to ∼50% of the entire population in Δ*mumT* strain, while only ∼5% of WT T6-cells were positive under oxidative stress supplemented with MnCl_2_ (Fig 2E and S4G). Taken together, these data suggest that deletion of MumT results in an increase in T6SS expression and thus MumT seems to have a negative impact on T6SS regulation in *A. baumannii* under oxidative stress (scenario 2).

**Figure 2.**
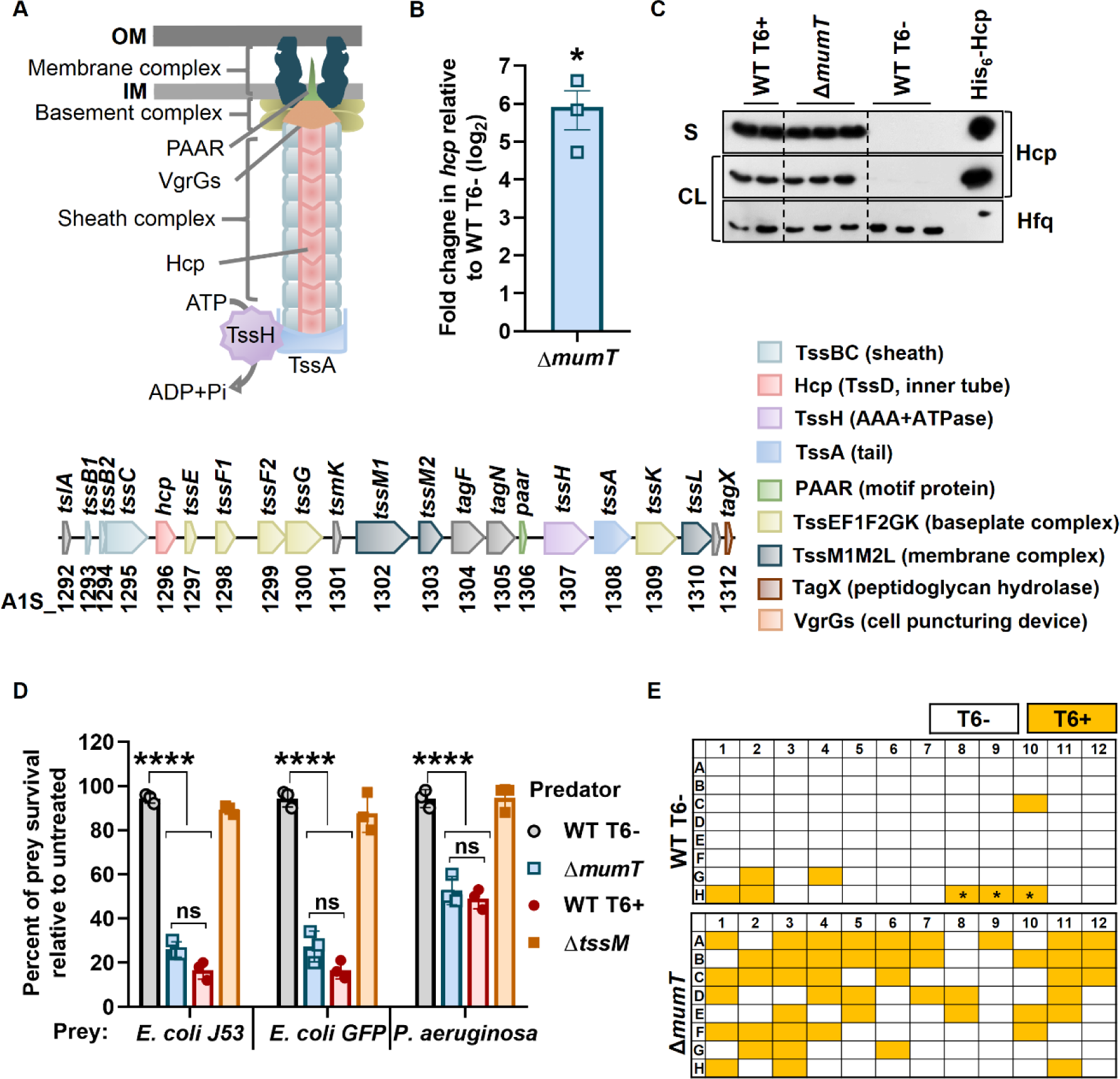
Deletion of *mumT* exhibited a significant increase in *hcp* expression under oxidative stress. (**A**) A schematic representation of T6SS (upper panel) and a schematic layout of the T6SS gene cluster in *A. baumannii* ATCC 17978 (lower panel). (**B**) Transcription of *hcp* was determined by qRT-PCR. The data represent the mean of three independent experiments, each in technical triplicates ± SEM. Statistical significance was determined using the Student’s t-test. * denotes p-value <0.05. (**C**) The Hcp secretion profile of the indicated strains was confirmed by a Western blot of cell lysate (CL) and culture supernatants (S). WT T6+ cell and purified His_6_-Hcp were used as a positive control for this assay. The Hfq antibody was used as a loading control for CL. The data represents three independent experiments where the samples were pooled and run on one gel. (**D**) In the T6SS competition assay, prey cells were subjected to killing by incubation with WT T6-, WT T6+, Δ*mumT*, and Δ*tssM* cells. The survival percentage of prey cells was enumerated by accounting for the respective untreated control (without predator/killer cells) as 100%. WT T6+ and Δ*tssM* (the deletion mutant of TssM in WT-17978, which does not have functional T6SS) cells were used as T6+ and T6-, respectively, control for this assay. The data represents the mean of three biological replicates ± SD. Statistical significance was determined using two-way ANOVA with Tukey’s multiple comparison test for each sample with the same prey killed by the killer strains. **** denotes p-value <0.0001. (**E**) Detection of Hcp secretion from individual colonies of WT T6-, and Δ*mumT* strains by Hcp-ELISA. * Well indicates purified His_6_-Hcp, used as a positive control for Hcp-ELISA.

### AbsR28 mediates the crosstalk between *mumT* and T6SS

We wondered how a metal transporter could impact the bacterial secretion system. Our observation that Δ*mumT* cells are predominant in T6+ despite having *tetR1* and *tetR2* (Fig S5A), which are transcriptional repressors of T6SS, suggests the presence of a MumT-dependent regulation of T6SS in *A. baumannii*. As the deletion of *mumT* in *A. baumannii* mimics an Mn^2+^-depleted stress condition (11), we speculated that the deletion of *mumT* might disrupt some transcriptional or post-transcriptional regulation of T6SS that causes T6SS upregulation in the Δ*mumT* cells. To begin with this hypothesis, we first checked the expression of several transcriptional regulators (i.e., *oxyR*, *zur*, and *fur*), which are known to regulate T6SS in other bacteria (13, 42, 43) at the transcriptional level in WT T6-, WT T6+, and Δ*mumT* cells grown under oxidative stress supplemented with MnCl_2_ using qRT-PCR. We also checked the transcript level of *mumR*, a transcriptional regulator of *mumT* in *A. baumannii* (12). None of the tested transcriptional regulators displayed any significant changes in expression (Fig S5B). sRNAs are known to play a vital role in post-transcriptional gene regulation in bacteria. The sRNA-mediated post-transcriptional regulation of T6SS is observed in *P. aeruginosa* (44–46). In light of this, we performed a CopraRNA search for all thirty-one putative sRNAs in *A. baumannii* that were predicted earlier using bioinformatic analysis by our lab (47), and eighteen of these sRNAs showed T6SS as a putative target (Table S1). To test the hypothesis further, we first picked four sRNAs (AbsR1, AbsR11, AbsR25, and AbsR28) out of those eighteen whose presence was already confirmed in *A. baumannii* and whose sequences were RACE mapped in our earlier studies (47). We checked the expression of four validated sRNAs (i.e., AbsR1, AbsR11, AbsR25, and AbsR28), and AbsR29 (which was not validated earlier to characterize it for its function), and *hfq* at the transcriptional level in WT T6-, WT T6+, and Δ*mumT* cells grown under oxidative stress supplemented with MnCl_2_ using qRT-PCR. Intriguingly, the expression of one sRNA, AbsR28, showed a notable fold reduction in the WT T6+ and Δ*mumT* cells compared with WT T6-cells (Fig 3A and S5C). Due to the maximum fold increase at the transcription level in the presence of MnCl_2_ under oxidative stress (Fig S5D) and significant fold reduction in WT T6+ cells and Δ*mumT* cells which are defective in Mn^2+^ utilization, we focused our study on evaluating the role of AbsR28 in *A. baumannii*’s T6SS regulation. We created a deletion mutant of AbsR28 in *A. baumannii* containing pAB3 plasmid (referred as ΔAbsR28 throughout the study) and confirmed that the deletion had no polar effect on transcript abundance (Fig S5E, F). To identify the genes regulated by AbsR28, we performed RNA-seq analysis comparing the relative abundance of total mRNA transcripts of WT T6- and ΔAbsR28 cells (upon exposure to oxidative stress and supplemented with MnCl_2_). In a comparison of WT T6- and ΔAbsR28 strains, the expression of seven structural genes of the T6SS [*tslA* (A1S_1292); *hcp* (A1S_1296); *tssF1* (A1S_1298); *tssF2* (A1S_1299); *tssG* (A1S_1300); *tsmK* (1301) and *tsgF* (A1S_1304)] and six genes encoding VgrGs/T6SS effector molecules (A1S_0086, A1S_0550, A1S_0551, A1S_1290, and A1S_3363) displayed a significant fold-induction in ΔAbsR28 strain (Fig 3B). All the genes encoding T6SS displayed a significant fold-increase in ΔAbsR28 cells compared to WT T6-cells grown under oxidative stress supplemented with MnCl_2_, indicating that AbsR28 negatively regulates T6SS (Fig. 3C). Like Δ*mumT* cells, ΔAbsR28 cells are also predominant in T6+ despite having *tetR1* and *tetR2* (Fig S5G). Next, we investigated whether AbsR28 regulates T6SS in *A. baumannii* and if there is a role of Mn^2+^ in this regulation. To evaluate this, we cloned AbsR28 under the control of an arabinose inducible promoter in the pWBAD30 vector and complemented the ΔAbsR28 strain. We observed that the elevated level of AbsR28 due to arabinose pulse, decreased the expression of Hcp, and the expression was further reduced when grown in the presence of MnCl_2_ in ΔAbsR28-pWBAD30AbsR28 strain (Fig 3D). The data was further validated by Hcp-western blot to detect secreted Hcp and a prey-predator assay, where increased expression of AbsR28 resulted in a notable reduction in Hcp secretion and increase in prey cell survival, respectively, (Fig 3E, F). These data indicate that AbsR28 efficiently represses T6SS expression in *A. baumannii* under oxidative stress in the presence of Mn^2+^.

**Figure 3.**
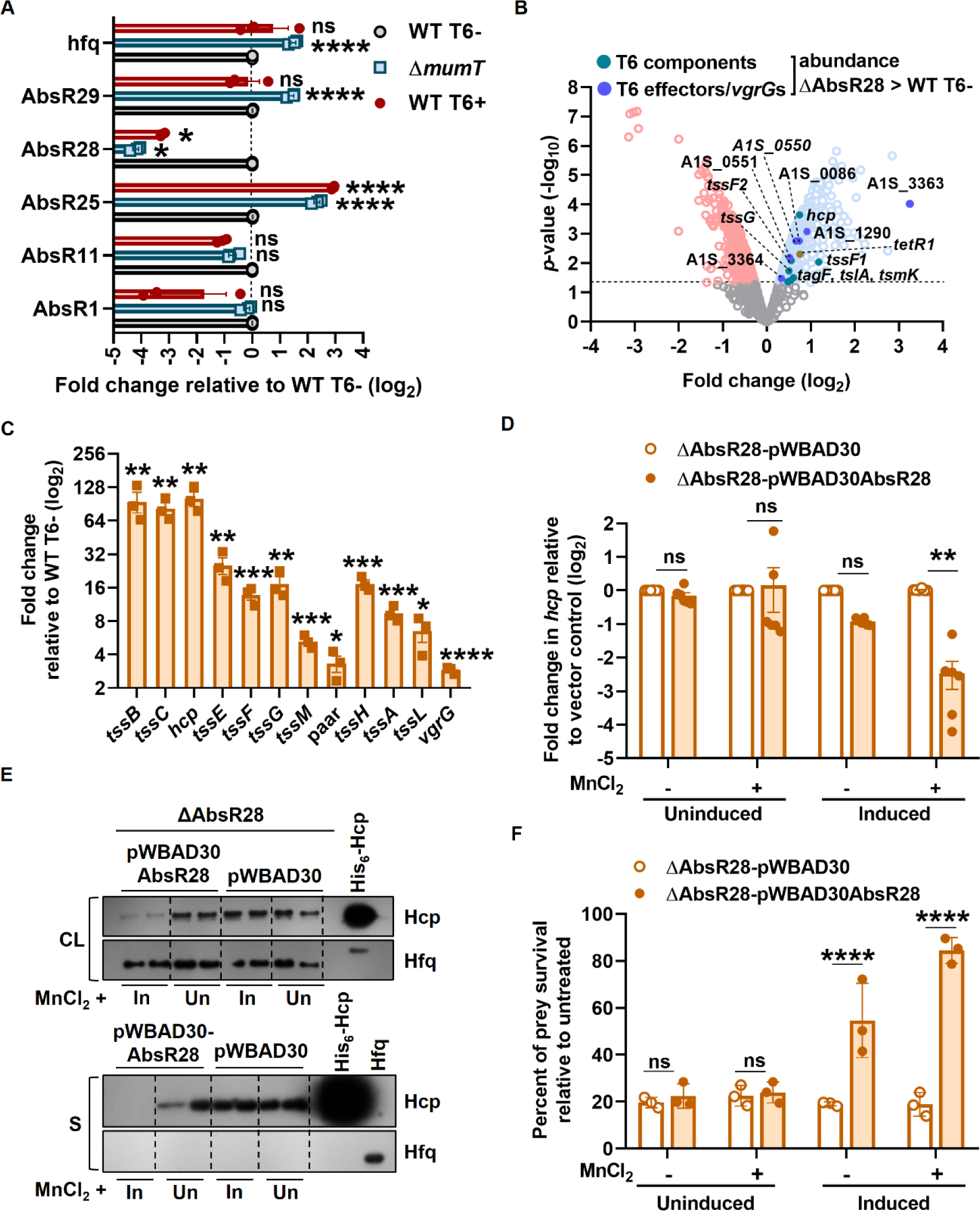
AbsR28 effectively represses T6SS in the presence of Mn^2+^. (**A**) The expression of several sRNAs and *hfq* was examined by qRT-PCR. The data represent the mean of three independent experiments, each in technical triplicates ± SEM. (**B**) RNA-seq analysis comparing RNA from MV and MnCl_2_ (both at 250 µM final conc.) treated ΔAbsR28 to a WT T6-control in duplicates. A dotted black line denotes p <0.05. (**C**) The transcription of each T6SS structural gene in WT T6- and ΔAbsR28 cells was determined by qRT-PCR. The data represent the mean of three independent experiments, each in technical triplicates ± SEM. (**D**) Transcription of *hcp* was determined by qRT-PCR. Un and In denote uninduced (without arabinose) and induced (with arabinose), respectively. The data represent the mean of three independent experiments, each in technical triplicates ± SEM. (**E**) The Hcp secretion profile of the bacterial strains (in biological duplicates) was tested by Western blot of cell lysate (CL) and cell-free supernatant (S). The Hfq antibody was used as a loading control for CL and to confirm that the supernatants were free from the bacterial cell. Un and In denote uninduced (without arabinose) and induced (with arabinose), respectively. (**F**) In the T6SS competition assay, *E. coli* J53 prey cells were subjected to killing by ΔAbsR28-pWBAD30AbsR28 and ΔAbsR28-pWBAD30 strains. The data represents the mean of three biological replicates ± SD.

### Binding of Mn^2+^ to AbsR28 alters its native structure

An sRNA’s ability to target a certain mRNA is contingent on its ability to form a base-pairing interaction with that mRNA at a specific location, known as the seed region (48). In AbsR28, the region that forms the most frequent base pairs with the predicted targets is positioned ∼1-50 nt from the start site, making it inaccessible for interaction due to the stem formation (Fig. 4A). Due to this, we hypothesized that Mn^2+^ might bind to AbsR28 and alter its native structure, which might be essential for base pairing and its regulatory function. The dissociation constant (K_D_) between Mn^2+^ and AbsR28 was 5.95×10^-6^ ± 1.21×10^-6^ M, as measured by isothermal titration calorimetry (ITC) (Fig 4B). The affinity to Mn^2+^ is more efficient than the controls where the binding of another small RNA (i.e., AbsR25) with Mn^2+^ and AbsR28 with Mg^2+^ were assessed (Fig S6A, B). Mutations in AbsR28 (U_89_, A_91_U_94_A_95_, and U_84_U_85_) reduced its affinity to Mn^2+^ except one mutant (A_76_A_77_) (Fig S6C). To determine whether the binding of Mn^2+^ affects the native structure of AbsR28, we performed *in vitro* structural probing with 5’-end labeled [γ-^32^P]ATP AbsR28 using lead(II) acetate (PbAc). The Mn^2+^ concentration-dependent changes in the cleavage of AbsR28 were observed in a denaturing-PAGE (Fig 4C), which were not very evident in the presence of Mg^2+^, used as a control (Fig 4D). These data confirm that Mn^2+^ binds to AbsR28 and alters its native structure.

**Figure 4.**
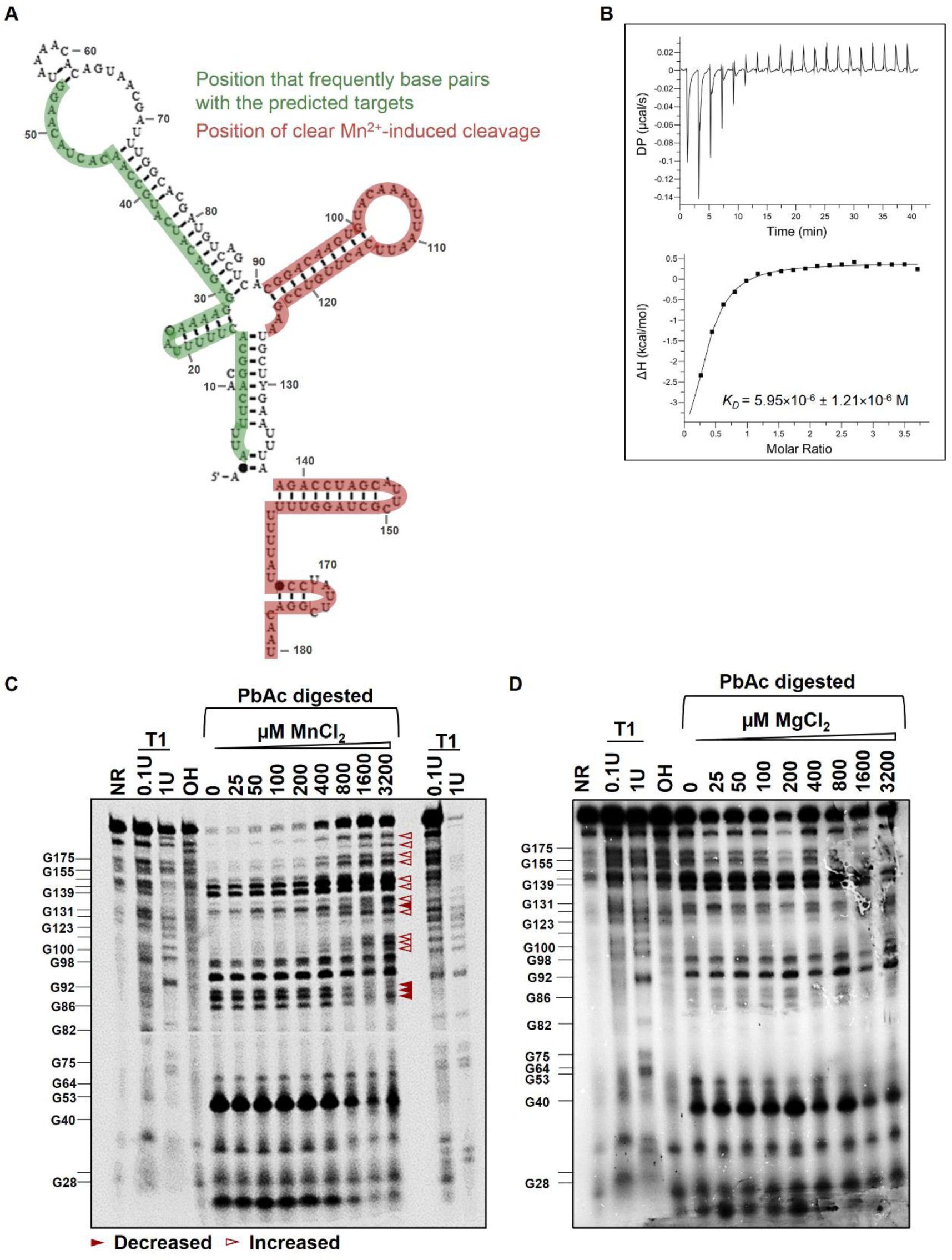
Mn^2+^ binds to AbsR28 and alters its structure. (**A**) The secondary structure of *A. baumannii* AbsR28 sRNA was predicted by Rfam software. (**B**) Isothermal titration calorimetry (ITC) of Mn^2+^ to AbsR28. (**C and D**) Lead acetate probing of the 5’end-labeled [γ-^32^P]ATP AbsR28 in an increasing concentration of either MnCl_2_ or MgCl_2_ provided in the structure buffer. Lanes indicated as T1 and OH ladders were obtained from the same labeled AbsR28 after incubation with RNase T1 and hydroxyl anions, respectively. RNase T1 digestion was performed in duplicates at 0.1 and 1.0 U concentrations. The position of cleaved G residues is marked at the left of the gel. Statistical significance was determined using the multiple comparison two-way ANOVA test with the Sidak correction for multiple comparisons comparing the means of each group to one another (A, D, F) and Student’s t-test (C). * denotes p-value <0.05, ** denotes p-value <0.01, *** denotes p-value <0.001, **** denotes <0.0001, ns denotes not significant.

### Mn^2+^ is required for AbsR28 to interact with *tssM* mRNA

sRNAs regulate gene expression by directly base pairing with target mRNA (49). Since T6SS in *A. baumannii* is a large gene cluster (50), we used the CopraRNA bioinformatic tool (51) to predict the targets of AbsR28 in *A. baumannii*. A CopraRNA search using the whole-genome sequence of *A. baumannii* ATCC 17978 and full-length AbsR28 (RACE mapped sequence) as inputs predicted base pairing between *tssM* mRNA position 258-291 and position 27-55 of AbsR28 with a predicted hybridization energy value of -14.10 kcal/mol (Fig 5A). To validate the direct interaction between AbsR28 and *tssM* mRNA, an *in vitro* RNA-RNA interaction was assessed. The addition of full-length AbsR28 at increasing concentrations to *tssM in vitro* transcripts resulted in the retardation of *tssM* transcripts in a native PAGE in the presence of Mn^2+^ (Fig 5B); in contrast, the retardation of *tssM* transcripts was very weak in the presence of Mg^2+^, used as a control (Fig 5C and 5D). The addition of full-length AbsR28 at increasing concentrations to an unrelated mRNA *pntB in vitro* transcripts (used as a negative control) did not show any retardation (Fig. S7A). For further validation of AbsR28-*tssM* binding, AbsR28 was mutated at the seed region (AbsR28-M9; A_27_G_28_G_29_A_30_ was mutated to U_27_C_28_C_29_U_30_) (Fig S6B). The retardation of *tssM* transcripts was much weaker even with higher concentrations of AbsR28-M9 compared to native AbsR28 (Fig S7C and S7D). Furthermore, we show that the compensatory mutation in *tssM* mRNA (*tssM*\*) restored the binding of AbsR28-M9 (S7E). Collectively, these findings indicate that the direct interaction of AbsR28 and *tssM* transcripts is Mn^2+^-dependent.

**Figure 5.**
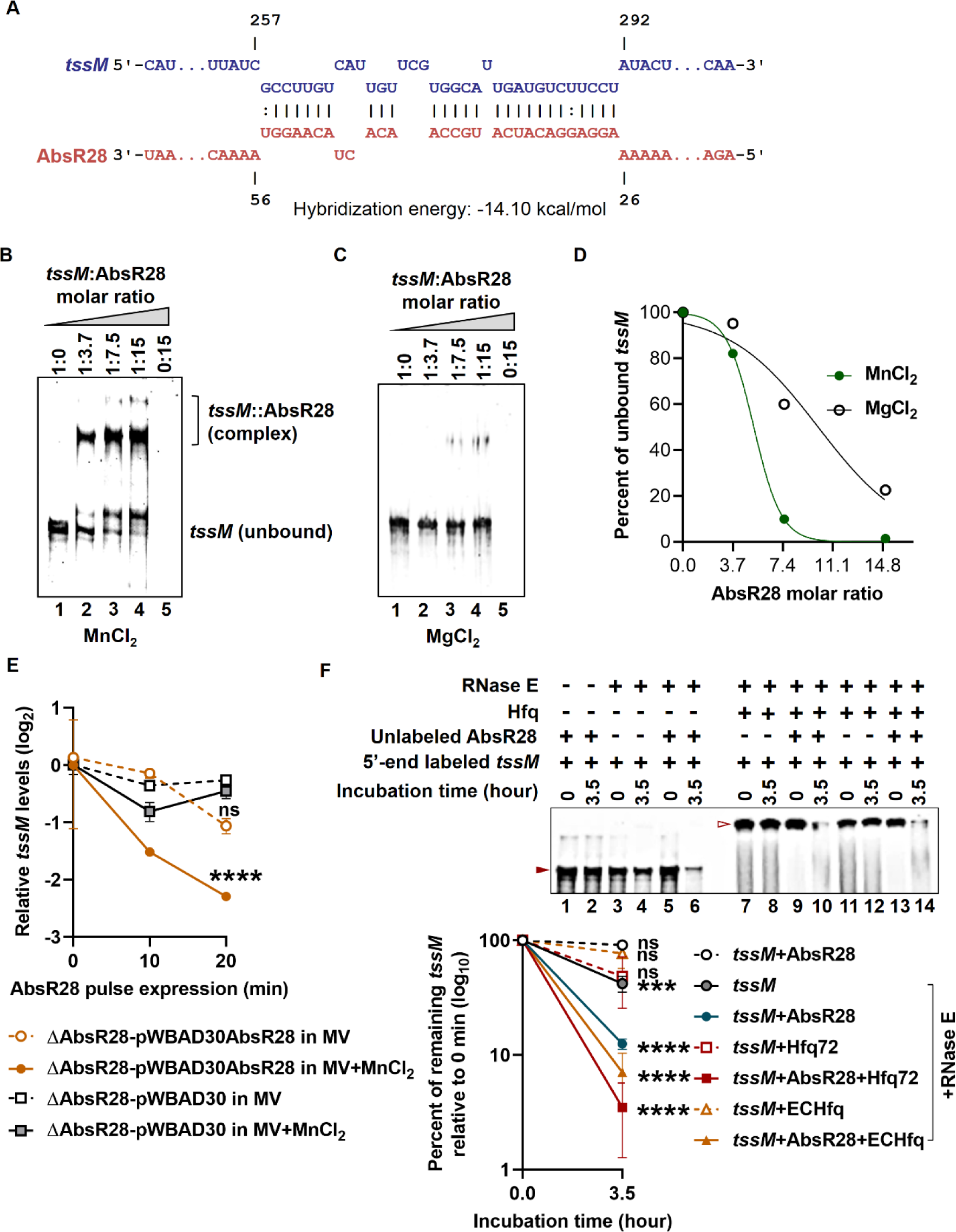
Mn^2+^ is required for AbsR28 to base pair with *tssM* mRNA and represses T6SS by potentiating RNase E-mediated degradation of *tssM* mRNA. (**A**) Predicted base pairing between *tssM* mRNA and AbsR28 by the CopraRNA bioinformatics tool. (**B and C**) Gel retardation assay of unlabeled *tssM in vitro* transcripts and unlabeled full-length AbsR28 *in vitro* transcripts in structure buffer containing either 10 mM MnCl_2_ or MgCl_2_. (**D**) Using ImageJ software, unbound *tssM* transcripts were quantified from Figures B and C (n = 2 independent experiments). Nonlinear regression was used to fit the curve. (**E**) The stability of *tssM* transcripts was determined after adding a RNA synthesis inhibitor Rifampicin (400 µg/mL), by qRT-PCR and plotted as the fold change relative to 0 min for each sample. The data represents the mean of technical triplicates ± SD. (**F**) The cleavage of the 5’end-labeled [γ-^32^P]ATP *tssM in vitro* transcripts was assessed by incubating with RNase E in structure buffer containing MnCl_2_ for 0 and 3.5 h in the presence of either unlabeled AbsR28 (lanes 5-6) or *A. baumannii* Hfq72 (lanes 7-8) or unlabeled AbsR28 + *A. baumannii* Hfq72 (lanes 9-10) or *E. coli* Hfq (lanes 11-12) or unlabeled AbsR28 + *E. coli* Hfq (lanes 13-14). Addition of Hfq results in a slower migration which is evident from the retardation of bands (7–14). The lower panel represents the quantification of the complete 5’-labeled *tssM* transcripts that remain after RNase E-mediated cleavage (denoted by a filled triangle from lanes 1-6 and an empty triangle from lanes 7-14) using ImageJ software (n = 3 independent experiments). Statistical significance was determined using the one-way ANOVA test with Dunnett’s multiple comparisons (E) and Student’s t-test (F). *** denotes p-value <0.001, **** denotes p-value <0.0001, ns denotes not significant.

### AbsR28 potentiates RNase E-mediated *tssM* mRNA degradation

To study the molecular mechanism of AbsR28-meditated T6SS repression in *A. baumannii*, we checked the effect of AbsR28-*tssM* interaction on the stability of target mRNA (i.e., *tssM*) via *in vivo* pulse expression. AbsR28 markedly reduced the stability of *tssM* mRNA *in vivo* in the presence of Mn^2+^ (Fig 5E). We observed that the elevated level of native AbsR28 reduced the level of *tssM* transcripts more significantly compared to the AbsR28-M9 via *in vivo* pulse expression assay (Fig S7F). This result confirmed that *tssM* processing depends on the direct interaction between AbsR28 and *tssM*. sRNAs induce the decay of target mRNA by RNase E-mediated mRNA degradation in an Hfq-dependent manner and repress its translation (52–58). We hypothesized that this might also happen in the case of AbsR28. To examine this, we performed an *in vitro* RNase E-mediated degradation assay where the 5’-end of the *in vitro* transcribed *tssM* was labeled with [γ-^32^P]ATP and incubated with purified RNase E in the presence or absence of unlabeled AbsR28 with *A. baumannii* Hfq72 or *E. coli* Hfq protein. RNase E alone cleaves *tssM* transcripts, but the cleavage is more prominent in the presence of AbsR28. As expected, there is a distinct RNase E-mediated degradation of *tssM* transcripts in the presence of AbsR28 and Hfq (Fig. 5F). These data demonstrate the molecular mechanism behind AbsR28-mediated T6SS regulation where the degradation of *tssM* mRNA by RNase E is strongly enhanced by base pairing to AbsR28 in the presence of Mn^2+^ and reduces the abundance of *tssM* transcripts for T6SS assembly.

### AbsR28 contributes to the fitness of *A. baumannii* during infection

In addition to T6SS regulation, we observed that the deletion of AbsR28 primarily affects genes involved in bacterial metabolism (carbohydrate, phenylalanine, cysteine, and methionine metabolism) and stress response (two-component system and ABC transporters) in RNAseq data (Table S2). Since AbsR28 remains uncharacterized in *A. baumannii* pathophysiology; therefore, we aimed to explore the importance of AbsR28-mediated gene regulation in the pathogenesis of *A. baumannii*. We co-incubated human blood-derived neutrophils with either WT T6-, WT T6+, Δ*mumT*, or ΔAbsR28 cells and checked their survival. WT T6-cells exhibited ∼85% survival, whereas ΔAbsR28 cells showed only ∼40% survival against neutrophil-mediated killing compared to untreated cells (Fig 6A). We next assessed the pathogenesis of WT T6-, WT T6+, Δ*mumT*, and ΔAbsR28 strains in a mouse model of *A. baumannii*-induced pneumonia where the strains were intranasally administered to BALB/c mice. In consistency with the previous observation that the deletion of *mumT* in *A. baumannii* results in compromised survival in mice due to a defect in Mn^2+^-uptake (12), we also observed that Δ*mumT* strain had a lower organ burden compared to the WT T6-strain. Additionally, WT T6+ and ΔAbsR28 strains had significantly reduced burdens in the lungs and liver when compared with the WT T6-strain (Fig 6B). Lungs isolated from mice infected with the WT T6-strain exhibited a remarkable tissue infiltration as indicated by necrosis and alveolar inflammation, whereas the mice lungs infected with either WT T6+, Δ*mumT*, or ΔAbsR28 strains did not show any notable damage (Fig 6C). Together, the data suggest that AbsR28 contributes to *A. baumannii* fitness during infection.

**Figure 6.**
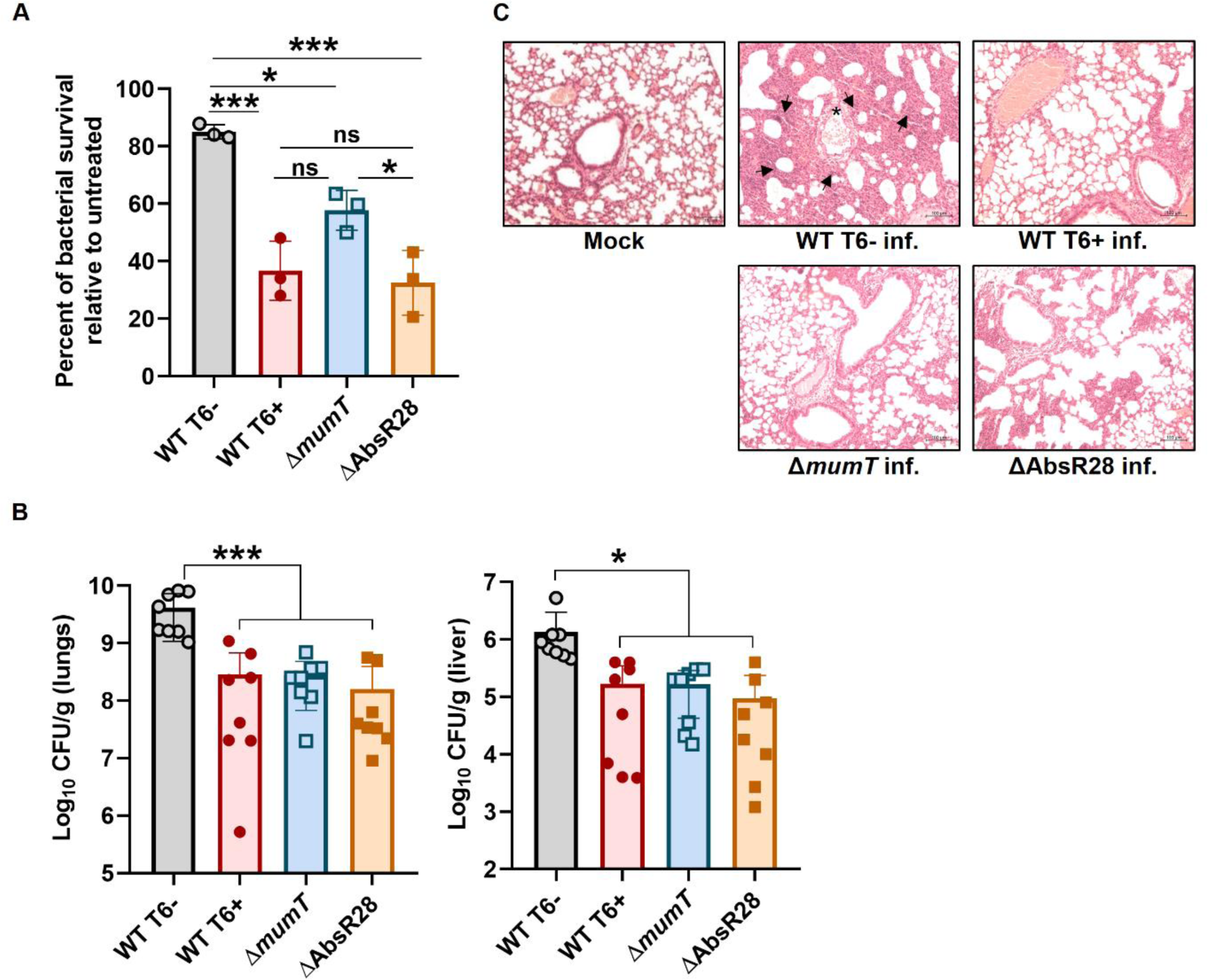
Mn^2+^-dependent AbsR28-mediated T6SS repression is required for *A. baumannii* pathogenesis. (**A**) Bacterial strains were co-incubated with neutrophils for 4 h at an MOI of 1:1. The percentage of bacterial survival was enumerated, accounting for the respective untreated control (without neutrophils) as 100%. The data represent the mean of three independent experiments, each in biological triplicates ± SEM. (**B**) Enumeration of bacterial burden recovered from mice lungs and liver (n = 8) infected with WT T6-, WT T6+, Δ*mumT*, or ΔAbsR28 strains at 36 hpi. Data represents mean ± SD. (**C**) Histopathology of the mice lungs (H&E stained) infected with WT T6-, WT T6+, Δ*mumT*, or ΔAbsR28 strains at 36 hpi. Tissue infiltration is indicated by necrosis (arrowheads) and alveolar inflammation (asterisks). The scale bar is 100 µM. Statistical significance was determined using the one-way ANOVA test with Tukey’s multiple comparisons (A, B). * denotes p-value <0.05, *** denotes p-value <0.001, ns denotes not significant.

## DISCUSSION

During bacterial infection, the host recruits phagocytic cells at the site of infection, which creates oxidative stress and limits the availability of free metal ions (12, 59, 60). For certain bacteria, the T6SS sequesters extracellular free metal ions and helps them cope with oxidative stress and host-mediated free metal restriction, so it is advantageous for them to express T6SS during oxidative stress (13, 44, 61). However, such role of T6SS in *A. baumannii*’s ability to survive under phagocytic cell-mediated oxidative stress has not yet been explored. Unlike *B. thailandensis*, where a synergistic relationship exists between the Mn^2+^-uptake system and T6SS expression (13), we found that *A. baumannii* cells that switch to T6+ cells are sensitive to oxidative stress due to inadequate intracellular Mn^2+^, which is required as a cofactor for SOD and catalase to break down intracellular ROS generated during oxidative stress (Fig 1 and S1). This distinguishes *A. baumannii* from other pathogens with distinctive gene regulatory mechanisms and emphasizes the need for studying *A. baumannii* pathophysiology in depth. To check the impact of intracellular Mn^2+^ on T6SS expression, we observed that the deletion of *mumT* promotes a significant increase in T6SS expression under oxidative stress (Fig 2). The T6SS expression is reduced due to adequate intracellular Mn^2+^ in the *mumT*-overexpressed strains (Fig S4), indicating the existence of a *mumT*-dependent T6SS regulation. Next, we focused on determining the molecular mechanism behind this regulation.

As deletion of *mumT* resulted in an increase in T6SS expression in *A. baumannii* (Fig 2) and there were no significant changes in the expression of known transcriptional regulators, we hypothesized that this might be due to Mn^2+^-dependent post-transcriptional repression, which is affected by the deletion of *mumT*. We observed that AbsR28 mediates the crosstalk between MumT and T6SS in *A. baumannii* during oxidative stress (Fig 3). Next, we focused on elucidating the mechanistic details of AbsR28-mediated T6SS repression. We show that the binding of Mn^2+^ to AbsR28 alters its native structure and results in a complementary base pairing with *tssM* transcripts (Fig. 4B, 4C and 5B). The exact Mn^2+^-binding region of AbsR28 remains to be determined. TssM is an indispensable structural component of the T6SS membrane complex which is essential for T6SS assembly (62–65). Hence, the deletion of *tssM* disrupts the functionality of T6SS in *A. baumannii* (66). We observed that the stability of *tssM* transcripts was significantly reduced when AbsR28 was pulse expressed in the presence of Mn^2+^ (Fig 5E). In Gammaproteobacteria, RNase E is the primary catalyst of mRNA decay (52). Since we observed AbsR28 expression-dependent decay of *tssM* transcripts *in vivo*, we hypothesized that AbsR28 triggers RNase E-mediated processing of *tssM* transcripts. Our in-line probing data demonstrate that AbsR28 potentiates RNase E-mediated degradation of *tssM* transcripts with the assistance of RNA chaperone Hfq (Fig 5F).

The T6SS is often not expressed in most of the clinical isolates of *A. baumannii* (67–69). The reason behind the inactive T6SS in *A. baumannii* is not fully understood. The following factors may account for potentially silent T6SS in *A. baumannii* clinical isolates: (i) high energy is required for T6SS expression, so to utilize the energy for other cellular processes, T6SS is silent; (ii) T6SS components are immunogenic (70, 71), so it may be advantageous for *A. baumannii* T6SS inactive strains to evade the host-mediated immune response (72); (iii) *A. baumannii* cells switch from T6-to T6+ at the expense of losing antibiotic resistance encoded by the plasmid (23); (iv) the T6+ cells of *A. baumannii* urinary tract infection (UTI) isolate have been previously reported to exhibit decreased virulence due to loss of pAB5 (a pAB3-family member), which regulates the expression of several chromosomally-encoded virulence factors (73); (v) firing the T6SS arrow involves bacteria rupturing the outer membrane, which could potentially disrupt the membrane integrity and render *A. baumannii* T6+ cells more sensitive to oxidative stress. Thus, to maintain the advantage of antibiotic resistance and virulence factors over the contact-dependent killing mechanism, it might be preferable for *A. baumannii* cells to inactivate the T6SS by other mechanisms independent of plasmid-mediated repression. A comparative study on the *A. baumannii* clinical strains and lab strains about the regulatory machinery of T6SS regulation during oxidative stress can be conducted in future for a better understanding of T6SS regulation.

In summary, we explored the molecular mechanism of AbsR28-mediated T6SS repression in *A. baumannii* ATCC 17978 (Fig 7). The sequential events of this proposed pathway are as follows: (i) during oxidative stress, *A. baumannii* cells upregulate *mumT* to increase Mn^2+^ uptake; (ii) intracellular Mn^2+^ binds to AbsR28 and alters its native structure; (iii) the altered structure of AbsR28 aids in the complementary base pairing with *tssM* mRNA transcripts and triggers RNase E processing; (iv) degradation by RNase E reduces the abundance of *tssM* transcripts and thereby represses the T6SS expression. *A. baumannii* cells that lose this controlled regulation result in T6+ phenotype. Our results unravel a novel mechanism of an Mn^2+^-dependent sRNA-mediated T6SS regulation in *A. baumannii*. Collectively, the findings presented here widen our knowledge about sRNA-mediated gene regulation and its role in shaping *A. baumannii*’s pathophysiology.

**Figure 7.**
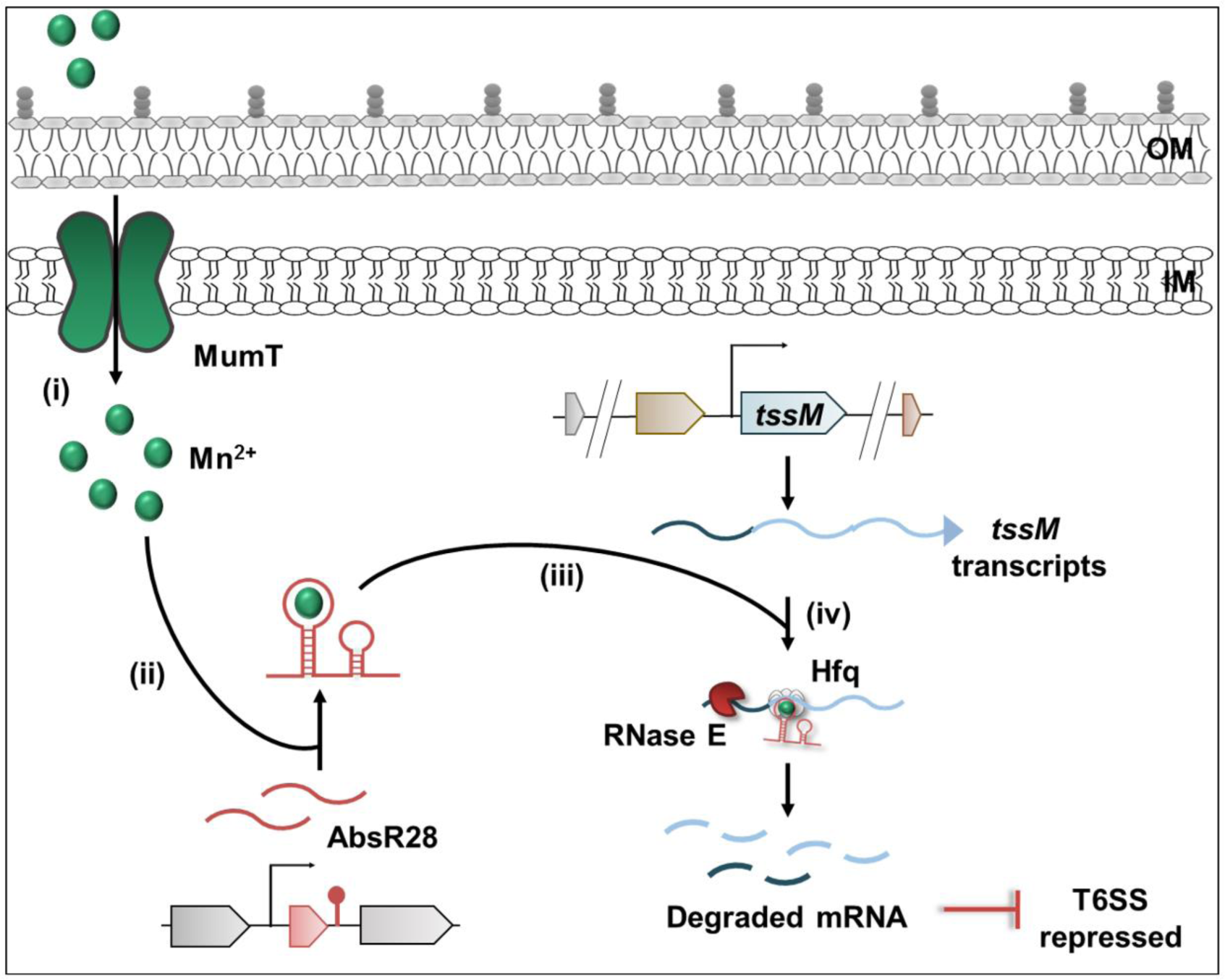
A proposed model of AbsR28-mediated post-transcriptional repression of T6SS in *A. baumannii* during oxidative stress. In the presence of Mn^2+^, AbsR28 base pairs with *tssM* mRNA and induces RNase E-mediated processing of *tssM*, which results in the repression of T6SS in *A. baumannii*.

## MATERIALS AND METHODS

### Bacterial strains and growth conditions

Bacterial strains and plasmids used in this study are listed in Tables S3 and S4. *A. baumannii* strains were grown at 37 ⁰C in LB broth or LB agar. *E. coli* J53 strains were grown in LB broth or LB agar, supplemented with 100 µg/mL sodium azide when necessary. *E. coli*-pNYL GFP strains were grown in LB broth or LB agar, supplemented with 50 µg/mL kanamycin when necessary. *P. aeruginosa* PAO1 strains were grown in LB broth or LB agar. *A. baumannii* strains that contain the pWBAD30 vector were grown in LB broth or LB agar, supplemented with 50 µg/mL kanamycin.

### Estimation of cell survival from phagocytosis

All experiments using human blood-derived neutrophils under protocol BT/IHEC-612020/7865 were reviewed and approved by the Institute Human Ethics Committee (HEC) of the Indian Institute of Technology Roorkee. The estimation of bacterial survival from phagocytic cell-mediated killing was performed as described previously (74). The detailed procedure is provided in the Supplemental material.

### Growth assay under oxidative stress

From the mid-log phase cultures of the indicated strains, 0.1% inoculum was inoculated in a 200 µL of fresh LB medium containing methyl viologen (MV) at 250 µM (final conc.). All the growth assays were performed in a sterile 96-well plate (Genaxy) at 37 ⁰C with shaking linearly at 180 CPM (6 mm), and the OD_600_ as a measurement of growth was measured at every 30 min interval for the indicated total time in the Synergy microplate reader (BioTek). Only media without any culture served as a negative control for this assay. The represented data is after background correction.

### ROS quantification

Bacterial cells were grown at 37 ⁰C with shaking to an OD_600_ of 0.6 (mid-log phase) in LB-medium. Bacterial cells were harvested by centrifugation and washed in sterile 1X PBS. The bacterial cell pellets were resuspended in 1X PBS, and 2’,7’-dichlorofluorescein diacetate (Thermo Fisher Scientific, D399) was added at a final concentration of 100 µM. After incubation for 30 min at 37 ⁰C, the cells were washed with 1X PBS to remove excess dye and transferred to 100 µL/well of a 96-well transparent bottom black well plate (BRAND). MV (250 µM final conc.) was added to the wells containing bacterial cells and incubated at 37 ⁰C with shaking linearly at 180 CPM (6 mm). OD_600_ and fluorescence (excitation/emission at 485/535 nm) were measured every 10 min interval for the indicated total time in the Synergy microplate reader (BioTek). The represented data is after background correction and OD_600_ normalization.

### Quantitative RT-PCR analysis

Human blood-derived neutrophils were co-incubated with the bacterial strains as described in the above section. After 4 h of incubation, the tissue culture plate was centrifuged at 400g for 5 min to settle down the neutrophils. Sample supernatants containing the bacterial cells were collected and harvested the bacterial cells by centrifugation at maximum speed. After washing the cell pellet with 1X PBS, RNA was extracted from the bacterial cells by the classic phenol-chloroform method. The cDNA synthesis was performed using PrimeScript 1st strand cDNA Synthesis Kit (TakaRa, 28704) according to the manufacturer’s instructions. Amplifications were achieved using a 3-step program on a QuantStudio 5 system (Thermo Fisher Scientific). Transcript abundance was calculated using the ΔΔC_T_ method (75) and normalized by the 16s gene.

### Quantification of intracellular metal content

The concentration of metal content in bacterial cells was determined as described previously (12) with some modifications. The detailed procedure is provided in the Supplemental material.

### Bacterial survival assay upon uptake of metal ions under oxidative stress

Bacterial cells were grown in 5 mL M9-medium (supplemented with 1% casamino acids) with or without MnCl_2_, ZnSO_4_, or FeCl_3_ at a final concentration of 250 µM and grown at 37 ⁰C with shaking to an OD_600_ of 0.6 (mid-log phase). The cultures were then treated with MV at a final concentration of 250 µM to induce oxidative stress and grow for another 2 h at 37 ⁰C with shaking. Cells were harvested from a 1 mL culture, washed with 1X PBS, and dissolved into 50 µL of 1X PBS. For spot assay, after a serial dilution in 1X PBS, 5 µL from each dilution was spotted onto LB agar containing MV at a final concentration of 100 µM, incubated at 37 ⁰C for O/N, and images were taken by a gel documentation system (Bio-Rad).

### Western blot analysis for Hcp

From the mid-log phase cultures of the indicated strains, 0.1% inoculum from the O/N cultures was then subcultured into fresh 5 mL of LB-medium containing MnCl_2_ at a final concentration of 250 µM and grown at 37 ⁰C with shaking to an OD_600_ of 0.6 (mid-log phase) for each strain. MV was added to the culture at a final concentration of 250 µM and incubated for another 2 h. The Hcp secretion phenotype of the strains was performed by Hcp-Western blot as described in Supplemental material. Primary anti-Hfq-antibody raised in rabbit at a dilution of 1:1000 was used as a loading control for cell lysate (CL).

### Bacterial killing assay

Bacterial killing assays were performed as described previously (23). *A. baumannii* T6- and Δ*mumT* strains were used as killers/predators, while *E. coli* J53 or *E. coli*-pNYL GFP, or *P. aeruginosa* were used as prey for this assay. The detailed procedure is provided in the Supplemental material.

### ELISA assay for HCP

From the mid-log phase cultures of the indicated strains, 0.1% inoculum from the O/N cultures was then subcultured into fresh 5 mL of LB-medium containing MnCl_2_ at a final concentration of 250 µM and grown at 37 ⁰C with shaking to an OD_600_ of 0.6 (mid-log phase) for each strain. MV was added to the culture at a final concentration of 250 µM and incubated for another 2 h. The rest of the protocol for Hcp-ELISA was performed as mentioned in Supplemental material.

### RNA-sequencing data analysis

RNA was isolated from WT T6- and ΔAbsR28 cells grown in LB supplemented with MnCl_2_ (250 µM) to an OD_600_ of 0.6 (mid-log phase), treated with MV (250 µM) for 2 h, and purified as described above. RNA sequencing was performed by Biokart India Pvt. Ltd. (India) using the Illumina HiSeq 4000 platform (Illumina). Analysis was performed by Rockhopper v 2.0.3. Comparative and statistical analyses were performed using iGeak v1.0a using the reference *A. baumannii* ATCC 17978 genome (NCBI: CP000521.1).

### Pulse expression studies

An arabinose inducible vector, pWBAD30, was modified from the pBAD30 backbone for this study. The pWBAD30-AbsR28 plasmid was transformed into ΔAbsR28 electrocompetent cells and the transformants were selected on LB-agar plates containing kanamycin (50 µg/mL). ΔAbsR28 transformed with pWBAD30 served as a vector control. The bacterial survival assay was performed as described Supplemental material in the presence of arabinose (0.2% w/v final concentration).

### Isothermal calorimetry

ITC titrations were performed using the ITC-200 (GE Healthcare). After degassing, 600 µM MnCl_2_ was placed into the syringe and 30 µM *in vitro* transcribed AbsR28 was placed in the reaction cell. ITC was performed over 20 injections, each 1.8 µl of MnCl_2_ with an interval of 120 s to allow for equilibration of the mixture between injections at a constant stirring of 500 rpm. All reactions were performed in a buffer containing 10 mM Tris, 100 mM KCl, and pH 8.0 at 25 ⁰C. A control experiment was performed at the same condition in the absence of sRNA. Analysis after subtracting the control data was performed using the Origin version 7.0 software provided with the system and data was fitted as an independent binding model.

### *In vitro* RNA transcription and 5’-end labeling

*In vitro* transcription (IVT) was performed as described earlier (76). The template for IVT was obtained using *A. baumannii* ATCC 17978 genomic DNA as a PCR template and forward primers containing the T7 promoter listed in Table S5. The detailed procedure is provided in the Supplemental material.

### *In vitro* structural probing

*In vitro* structural probing was performed as described earlier (77) with some modifications using *in vitro* transcribed AbsR28 5’-labelled with [^32^P]-γ-ATP. The detailed procedure is provided in the Supplemental material.

### Gel retardation assay

The gel retardation assay was performed as described earlier (78) with some modifications using unlabeled *tssM in vitro* transcripts (250 nt upstream and 250 nt downstream from ATG) and full-length AbsR28 *in vitro* transcripts. The detailed procedure is provided in the Supplemental material.

### RNase E-mediated degradation assay

Rnase E-mediated degradation assay was performed using the *in vitro* transcript *tssM* 5’-labeled with [^32^P]-γ-ATP. The detailed procedure is provided in the Supplemental material.

### Mice infection model for *A. baumannii* pneumonia

All animal experiments under protocol BT/IAEC/2018/07 were reviewed and approved by the Institute Animal Ethics Committee of the Indian Institute of Technology Roorkee. Adult (6–8 week old) age-matched male BALB/c mice (n = 8 for each group, determined using G*Power analysis) were infected intranasally with *A. baumannii* and CFU was enumerated as previously described (79). Briefly, the mice were anesthetized and infected intranasally with 20 µL of inoculum containing 4×10^4^ CFU of the indicated strains. Mice were euthanized at 36 h of infection, and the lungs and livers were harvested, immediately transferred on ice, and washed with ice-cold 1X sterile PBS. The harvested organs were chopped into pieces to enumerate the bacterial burden, histopathology, and Western blot analysis.

### Quantification and statistical analysis

Statistical analyses were performed using GraphPad Prism 8. Quantifying bands from gels was performed using ImageJ and analyzed using GraphPad Prism 8. The details about the number of repeated experiments, statistical tests used, significance values, and group sizes are indicated in each figure legend.

## ACKNOWLEDGEMENTS

We thank all the members of the Pathania Lab for their suggestions and fruitful discussions regarding this manuscript. We are grateful to Prof. S.P. Mukherjee for the training required to handle [γ-^32^P]ATP at IIT Roorkee. We extend heartfelt thanks to Prof. N.K. Navani (IIT Roorkee) for his scientific inputs during experiments and critical views on the manuscript. We are thankful to Dr. Arpita Dey for helping with the ITC assay and data analysis. We thank Dr. Amit Gaurav for helping with RNA sequencing data deposition on the NCBI SRA. We thank Prof. P. Roy (IIT Roorkee) and members of his laboratory for animal cell lines. We also thank Dr. Siva for the clonings required for antibody generation and construction of TssM knockout in ATCC 17978. We are grateful to the Institute Instrumentation Centre (IIC) in IIT Roorkee for providing facilities like ICP-MS and ITC. This work was supported by the DBT/Wellcome Trust India Alliance Fellowship (grant number IA/S/21/1/505588) awarded to R.P. S.B. was supported by a Research Fellowship (Sr. No. 2061530396; Ref. No. 21/06/2015(i)EU-V) from University Grants Commission (UGC). A.P. was supported by a fellowship (Ref. No. 09/143(1019)/2020-EMR-I) from CSIR. K.D. was supported by a fellowship (Ref. No. DST/INSPIRE/03/2017/000008) from DST. A.S. was supported by a fellowship from MHRD. S.P. was supported by a fellowship (Ref. No. DBTHRDPMU/JRF/BET-22/I/2022-23/196) from DBT. The funders had no role in study design, data collection and interpretation, or the decision to submit the work for publication.

## AUTHOR CONTRIBUTIONS

S.B. and R.P. designed the study, conceptualized the experiments and interpreted the data. S.B. performed most of the experiments, analyzed the data, and prepared figures. A.P., K.D., A.S., S.P., and N.J. helped during the experimentation. T.B. generated preliminary data for AbsR28 that implied its role in oxidative stress. J.A. helped during animal experiments. R.S. S.C. and T.K.S. provided the radioactive facility for lead acetate probing at THSTI. R.P. engaged in the acquisition of funds. S.B. and R.P. wrote the manuscript,

R.P. edited the manuscript and all the authors commented on it.

## DATA AVAILABILITY

Raw RNA-sequence data have been deposited at NCBI Sequence Read Archive (NCBI SRA) with accession number PRJNA865908 and are publicly available as of the date of publication. Target prediction of AbsR28 was performed using CopraRNA (https://rna.informatik.uni-freiburg.de/CopraRNA [accessed 2nd October 2020]) using the RACE-mapped sequence of AbsR28 (45) as input for *Acinetobacter baumannii* ATCC 17978 (NCBI Reference Sequence: NZ_CP018664, NZ_CP049363, NZ_CP039028, and NZ_CP039023) as an organism of interest. Any additional information required to analyze the data reported in this paper is available upon request from the lead contact.

