## Supplemental material for "*Acinetobacter baumannii* represses type VI secretion system through an Mn^2+^-dependent sRNA-mediated regulation"

### DETAILED METHODS

#### Isolation of *A. baumannii* T6- and T6+ cells

*A. baumannii* T6- and T6+ cells were isolated using Hcp-ELISA as described previously (1) with some modifications. Briefly, a fresh colony of wild-type *A. baumannii* ATCC 17978 streaked on LB-agar plate was used to inoculate in 5 mL of LB-medium overnight (O/N) at 37 $^0$C with shaking. 0.1% inoculum from the O/N culture was then subcultured into fresh 5 mL of LB-medium and grown at 37 $^0$C with shaking to an $OD_{600}$ of 0.6 (mid-log phase). The culture was serially diluted in LB medium and plated on LB-agar, followed by incubation at 37 $^0$C for O/N to obtain more than 100 isolated colonies. Individual colonies were inoculated in a 96-well plate containing 200 µL of LB medium/well and incubated at 37 $^0$C for O/N with gentle shaking. After O/N growth, the plate was centrifuged to pellet down the bacterial cells and 75 µL of the supernatant from each well was transferred to a 96-well ELISA plate (Thermo Fisher Scientific) containing 25 µL of binding buffer (0.0258 M sodium carbonate, 0.0742 M sodium bicarbonate, pH 9.5) in each well (for example, A1 supernatant from the 96-well plate was transferred to A1 of 96-well ELISA plate). The ELISA plate was incubated at 4 $^0$C for O/N on a gel rocker for efficient binding. The plate was then washed with 1X PBS and blocked with 200 µL of blocking solution (5% w/v skim milk in PBST; 1X PBS containing 0.1% inoculum v/v Tween-20) for 1 h at room temperature (RT). Primary anti-Hcp-antibody raised in the rabbit at a dilution of 1:10000 in blocking solution (2.5% w/v skim milk in PBST) was used at 100 µL/well to probe at 4 $^0$C for O/N. Following three successive washes with PBST, HRP-conjugated goat anti-rabbit secondary antibody (Thermo Fisher Scientific, 31460) at a dilution of 1:20000 in a solution (1X PBS containing 0.1% inoculum v/v Tween-20) was added 100 µL/well and incubated for 1 h at RT in the dark. After three successive washes with PBST and one wash with PBS, 50 µL substrate (citrate phosphate buffer at pH 5.6 containing $H_2O_2$ and *o*-Phenylenediamine dihydrochloride) was added to each well and waited for 10-15 min to develop a yellow color. The reaction was stopped by adding 3N HCl, which will turn yellow to orange, and measured the OD at 495 nm. Purified $His_6$-Hcp was used as a positive control, and unsupplemented LB medium as a negative control for the assay. A well that appears to have the T6SS+ signal (i.e., develops an orange color after ELISA) was marked and the cells from that particular well of the 96-well plate (the source of the supernatant sample) were isolated (considered at *A. baumannii* T6+ after 1$^{st}$ round of ELISA). Similarly, a well that appears to have the T6SS- signal (i.e., does not develop any color after ELISA) was marked and the cells from that particular well of the 96-well plate (the source of the supernatant sample) were isolated (considered at *A. baumannii* T6- after 1$^{st}$ round of ELISA). The *A. baumannii* T6- and T6+ cells from 1$^{st}$ round ELISA were plated on LB agar, and further Hcp-ELISA (2$^{nd}$ round) was performed using freshly isolated individual colonies to confirm the T6SS phenotype.

Further, *A. baumannii* T6+ cells were checked for Hcp secretion phenotype by Hcp-Western blot. The *A. baumannii* T6- and T6+ cells from the 2$^{nd}$ round of ELISA were inoculated in LB medium and grew overnight (O/N) at 37 $^{0}$C with shaking. Individually, 0.1% inoculum from the O/N cultures was subcultured into fresh 5 mL of LB-medium and grown at 37 $^{0}$C with shaking to an $OD_{600}$ of 0.6 (mid-log phase) for both strains. Bacterial cells were harvested from 1 mL cultures and supernatants were collected. The supernatants were filtered through a 0.22 µm syringe filter (Merk Millipore Ltd.) and concentrated using trichloroacetic acid. The pellets were dissolved in 1X SDS-gel loading dye and heated at

95 $^{0}C$ for 5 min. Whole-cell lysate ($OD_{600}$ normalized volume) and supernatants were run on a 15% SDS-PAGE for separation and transferred to a PVDF membrane (Cytiva, GE10600023). Following blocking (5% w/v skim milk in PBST; 1X PBS containing 0.1% inoculum v/v Tween-20) for 1 h at RT, the membrane was probed by primary anti-Hcp-antibody raised in the rabbit at a dilution of 1:1000 in blocking solution (2.5% w/v skim milk in PBST) at 4 $^{0}C$ for O/N on a gel rocker. Following five successive washes with PBST, HRP-conjugated goat anti-rabbit secondary antibody (Thermo Fisher Scientific, 31460) at a dilution of 1:20000 in a solution (1X PBS containing 0.1% inoculum v/v Tween-20) was added and incubated for 1 h at RT in the dark. After five successive washes with PBST and one wash with PBS, ECL substrate (TakaRa) was added and developed onto an X-ray film.

After isolating WT T6- and WT T6+ variants from the wild-type strain *A. baumannii* ATCC 17978 by Hcp-ELISA and validation by Hcp-Western blot, glycerol stocks were prepared. The WT T6- and WT T6+ variants were freshly streaked, and the phenotype was confirmed by several Hcp-ELISA assays. Before and after each experiment, the Hcp-secretion profiles of the WT T6- and WT T6+ strains were checked by Western blot to confirm the phenotypes.

**Estimation of cell survival from phagocytosis**

Neutrophils were isolated from human blood as described above and diluted to obtain a final concentration of $1x10^{4}$ cells/well. A freshly streaked colony of WT T6- and WT T6+ strain on LB-agar plates were used to inoculate in 5 mL of LB-medium for O/N at 37 $^{0}C$ with shaking. 0.1% inoculum from the O/N cultures was then subcultured into fresh 5 mL of LB-medium and grown at 37 $^{0}C$ with shaking to an $OD_{600}$ of 0.6 (mid-log phase).

Bacterial cells were harvested and diluted to obtain $10^4$ CFU/µL. The diluted bacterial cultures were then opsonized in fetal bovine serum (non-heat treated) for 15 min. Neutrophils were then co-incubated with bacterial strains at an MOI of 1:1 ratio in RPMI 1640 cell culture medium (HIMEDIA, AL028A) and incubated at 37 $^0$C in an animal tissue culture incubator (Eppendorf). The same method was performed for the macrophage RAW 264.7 cell line in the DMEM medium (HIMEDIA, AL007A). After 4 h of infection, the medium supernatants were serially diluted and plated onto Leeds *Acinetobacter* medium plates. After incubation at 37 $^0$C for O/N, the bacterial colonies were enumerated and the percent growth was quantified by dividing the CFU of the particular *A. baumannii* strain-neutrophil/macrophage RAW 264.7 cell line co-culture by that respective strain alone culture (grown in the same conditions in the absence of neutrophil/macrophage RAW 264.7 cell line). Only the neutrophil/ macrophage RAW 264.7 cell line was kept as a negative control for the assay.

**Transformation of pAB3 into WT T6+ cells**

Total plasmids (pAB1, pAB2, and pAB3 present in *A. baumannii* ATCC 17978 strain) were isolated from an overnight culture of *A. baumannii* ATCC 17978 strain grown in LB using plasmid miniprep kit (Thermo Fisher Scientific, K0503). The presence of pAB3 in the plasmid isolate was confirmed by PCR using *tetR1* and *tetR2* primers. The plasmid mixture was transformed into the electro-competent WT T6+ cell (devoid of pAB3) and the transformants were selected on LB agar plate containing sulfamethoxazole/trimethoprim (S&T; 30 µg/mL and 5 µg/mL, respectively). The transformation of pAB3 into WT T6+ competent cells was confirmed by PCR using forward and reverse primers of *tetR1* and *tetR2* genes.

**Generation of knockout strains**

The deletion mutants were created using a homologous-recombination method described previously (2) with some modifications. Briefly, a construct carrying an apramycin cassette (amplified from pMDIAI and having FRT sites on both sides) flanking between 500 bp upstream and 500 bp downstream of the gene of interest was cloned into a pUC18 vector (used as a cloning vector). A PCR product was amplified from the construct using a 125 bp upstream forward primer and a 125 bp downstream reverse primer of the gene of interest listed in Table S5. Around 5 µg of the concentrated PCR gel-purified product was transformed into *A. baumannii* electrocompetent cells harboring pAT02 (which contains $Rec_{Ab}$ system) under IPTG induction (2 mM) and plated on LB-agar containing apramycin (15 µg/mL). The transformants were further passaged on LB-agar containing an increasing concentration of apramycin (15-30 µg/mL). The recombinants were further confirmed by PCR using primers located outside the regions of homology (i.e., 500 bp upstream forward primer and 500 bp downstream reverse primer of the gene of interest) listed in Table S5. Following PCR confirmation and curing of pAT02, a clean knockout (K/O) was created by transforming pAT03 (which contains the FLP recombinase system) under IPTG induction (2 mM). A loss of apramycin resistance confirmed the clean K/O and the losing apramycin-FRT was further confirmed by PCR using apramycin forward and reverse primers listed in Table S5. After curing pAT03, the clean K/O strains were maintained in glycerol (15%) at -80 $^{0}$C for further use.

**Quantification of intracellular metal content**

A fresh streaked colony of WT T6- and WT T6+ strain on LB-agar plates were used to inoculate in 5 mL minimal medium (M9-medium) supplemented with 1% casamino acids as a nutrient source for O/N at 37 $^{0}$C with shaking. 0.1% inoculum from the O/N cultures was then subcultured into fresh 100 mL M9-medium (supplemented with 1% casamino acids) supplemented with or without $MnCl_2$, $ZnSO_4$, or $FeCl_3$ at a final concentration of 100 μM and grown at 37 $^{0}$C with shaking to an $OD_{600}$ of 0.6 (mid-log phase) for each strain. Then MV was added at a final concentration of 100 μM to the culture to induce oxidative stress, and it was grown further at 37 $^{0}$C with shaking for 4 h. The bacterial cultures ($OD_{600}$ normalized volume) were then transferred to pre-weighed metal-free 50 mL centrifuge tubes and centrifuged to harvest the cell pellet, washed thrice with Milli-Q deionized water, and dried thoroughly. The pellet weight was measured using an analytical balance (G&G). Pellets were digested with 1 mL of 70% $HNO_3$ using Milli-Q deionized water as a diluent for O/N at 90 $^{0}$C and diluted with 9 mL of 3.5% $HNO_3$ using Milli-Q deionized water as a diluent. The samples were then subjected to inductively coupled plasma-mass-spectrometry (8900 ICP-MS Triple Quad, Agilent) at the Institute Instrumentation Centre (IIC) in IIT Roorkee. The concentrations were determined by utilizing a standard curve for each metal. Only M9-medium supplemented with 1% casamino acids was used as a control.

**Bacterial killing assay**

*A. baumannii* T6SS- and Δ*mumT* strains were grown in 5 mL of LB-medium containing 100 μM MV and $MnCl_2$ at 37 $^{0}$C with shaking to an $OD_{600}$ of 0.6 (mid-log phase). Cells were harvested from a 2 mL culture, washed with 1X PBS, and dissolved into 50 μL of 1X PBS. Simultaneously, the prey cells were grown in LB medium containing respective selection markers to an $OD_{600}$ of 0.6 (mid-log phase), harvested the cells from 2 mL culture, washed with 1X PBS, and dissolved into 50 μL of 1X PBS. The predator and prey cells were mixed at a ratio of 1:1 and spotted 100 µL mixture on a sterile 0.22 µm syringe filter (Merk Millipore Ltd.) placed on dry LB agar plates. After air-drying inside the hood, the plates were kept at 37 $^{0}$C for 4 h. The mixed cultures were scraped out and resuspended in 1X PBS. For spot assay, after a serial dilution in 1X PBS, 5 µL from each dilution was spotted onto LB agar containing sodium azide (100 µg/mL) when *E. coli* J53 was used as prey or LB agar containing kanamycin (50 µg/mL) when *E. coli*-pNYL GFP was used as prey. The plates were incubated at 37 $^{0}$C for O/N and images were taken using a camera. For CFU count, after a serial dilution in 1X PBS, 100 µL from each dilution was spread onto LB agar containing sodium azide (100 µg/mL) when *E. coli* J53 was used as prey, or LB agar containing kanamycin (50 µg/mL) when *E. coli*-pNYL GFP was used as prey, or *P. aeruginosa* agar medium when *P. aeruginosa* was used as prey. The plates were incubated at 37 $^{0}$C for O/N. The survival percentage of the prey cells was calculated by considering the CFU of prey cells alone as 100%. To measure the prey cells' GFP fluorescence, the predator and prey cells were mixed at a 1:1 ratio in LB-medium in a 96-well transparent bottom black well plate (BRAND). GFP fluorescence was recorded at 485/525 nm to measure prey cells' growth at 37 $^{0}$C every 3 h.

**Pulse expression studies**

For survival assay, fresh colonies of the indicated strains streaked on LB-agar plates containing kanamycin (50 µg/mL) were used to inoculate in 5 mL of LB-medium containing kanamycin (50 µg/mL) or O/N at 37 $^{0}$C with shaking. 0.1% inoculum from the O/N cultures was then subcultured into fresh 5 mL of LB-medium with kanamycin (50 µg/mL) containing $MnCl_2$ at a final concentration of 250 µM and grown at 37 $^{0}$C with shaking to an $OD_{600}$ of 0.6 (mid-log phase). MV (250 µM final conc.) was added to the culture and grown for another 2 h. Cells were harvested from a 2 mL culture, washed with 1X PBS, and dissolved into 50 µL of 1X PBS. Simultaneously, *E. coli* J53 as prey cells were grown in LB medium containing sodium azide (100 µg/mL) to an $OD_{600}$ of 0.6 (mid-log phase), harvested the cells from 2 mL culture, washed with 1X PBS, and dissolved into 50 µL of 1X PBS. The predator and prey cells were mixed at a ratio of 1:1, arabinose (0.2% w/v final concentration) was added for AbsR28 expression and spotted the 100 µL mixture on a sterile 0.22 µm syringe filter (Merk Millipore Ltd.), placed on dry LB agar plates. After air-drying inside the hood, the plates were kept at 37 $^{0}$C for O/N. The mixed cultures were scraped out and resuspended in 1X PBS. After a serial dilution in 1X PBS, 100 µL from each dilution was spread onto LB agar containing sodium azide (100 µg/mL). The plates were incubated at 37 $^{0}$C for O/N. The survival percentage of the prey cells was calculated by considering the CFU of prey cells alone as 100%. To check gene expression by qRT-PCR, the cells were grown in LB containing $MnCl_2$ at a final concentration of 250 µM and grown at 37 $^{0}$C with shaking to an $OD_{600}$ of 0.6 (mid-log phase). MV (250 µM final conc.) and arabinose (0.2% w/v final concentration) were added to the media and incubated for 4 h. RNA was extracted, and qRT-PCR was performed as described above. For the Hcp-Western blot, the cells were grown in LB containing $MnCl_2$ at a final concentration of 250 µM and grown at 37 $^{0}$C with shaking to an $OD_{600}$ of 0.6 (mid-log phase). MV (250 µM final conc.) and arabinose (0.2% w/v final concentration) were added to the media and incubated for a further 4 h. Cell lysate (CL) and cell-free supernatant (S) were run on a SDS-PAGE and performed Western blot

##### *In vitro* RNA transcription and 5'-end labeling

T7 transcription was performed using the T7 RNA polymerase (Thermo Scientific, EP0111) according to the manufacturer's instructions. The DNA contamination was removed by incubating the transcript RNA with DNase I (Thermo Scientific, EN0521) according to the manufacturer's instructions. Complete transcripts were obtained by phenol-chloroform extraction, running on a long Urea-PAGE followed by gel purification and concentrated by sodium acetate precipitation. To perform 5'-end labeling, transcripts were dephosphorylated with FastAP (Thermo Scientific, EF0651) and 5'-labeled with [$^{32}$P]-γ-ATP using T4 polynucleotide kinase (Thermo Scientific, EK0032) with forward reaction buffer according to the manufacturer's protocol. Radiolabeled transcripts were purified by running a long Urea-PAGE followed by gel purification and concentrated by sodium acetate precipitation.

##### *In vitro* structural probing

*In vitro* transcribed 5'-labeled AbsR28 (2.5 pmol) was incubated in a modified 2X in-line buffer (100 mM Tris-HCl pH 8.3, 200 mM KCl) at a range of $MnCl_2$ concentrations containing yeast RNA (1 µg/reaction) for 40 h at room temperature. Afterward, 2 µL of 25 mM lead(II) acetate (Sigma-Aldrich, 215902) stock was added to each of the 10 µL reactions and incubated for precisely 2 min at 37 $^0$C. RNaseT1 ladder was generated by incubating 5'-labeled AbsR28 with RNaseT1 (0.1 U/µL and 1.0 U/µL) in 1X sequencing buffer (Ambion, AM2283) for 3 min at 55°C. Alkaline RNA ladders were generated by incubating 5'-labeled AbsR28 in 1X alkaline buffer (Ambion, AM2283) for 5 min at 90 $^0$C. All reactions were stopped immediately by adding a stop buffer (Ambion, AM2283). After phenol:chloroform:isoamyl alcohol purification, RNA pellets were dissolved in loading buffer II (Ambion, AM2283). All samples were denatured for 3 min at 95 $^0$C and loaded on 10% PAGE/7 M urea sequencing gels at a constant 15 Watt. After gel drying for 2 h at 80 $^0$C, bands were visualized using a phosphoimager (Typhoon FLA 7000, GE Healthcare) and ImageQuant software.

**Gel retardation assay**

Unlabeled *tssM in vitro* transcripts (250 nt upstream and 250 nt downstream from ATG) at a fixed concentration of 20 pmol and full-length AbsR28 *in vitro* transcripts at an increasing concentration were used for the gel retardation assay. The 10X structure buffer (100 mM Tris-HCl pH 7.0 and 1 M KCl) was added at a final concentration of 1X to the mRNA and the mRNA was allowed to re-nature at 37 $^0$C for 15 min. Yeast RNA (Ambion, AM2283) was added to the reaction mix at a final concentration of 1 µg/reaction and AbsR28 was added to the tubes at an increasing concentration. $MnCl_2$ was added to the reaction mix at a final concentration of 10 mM. After incubation at 37 $^0$C for 60 min, 6X RNA native loading buffer (Ambion) was added to stop the reaction and resolved on a 6% native PAGE at 4 $^0$C in 0.5% TBE at a constant current of 40 mA for 6 h. The gel was stained with SYBR Safe, and visualized the RNA bands using a phosphorimager (Typhoon FLA 9000, GE Healthcare) and ImageQuant software.

**RNase E-mediated degradation assay**

The *in vitro* transcript *tssM* (250 nt upstream and 250 nt downstream from ATG) was 5'-labeled with [$^{32}$P]-γ-ATP as described above. 5’-labeled *tssM* mRNA (5 pmol) was denatured at 65 $^0$C for 2 min and chilled on ice for 5 min. The 10X structure buffer (100 mM Tris-HCl pH 7.0 and 1 M KCl) was added at a final concentration of 1X to the mRNA and the mRNA was allowed to re-nature at 37 $^0$C for 15 min. Yeast RNA (Ambion,

AM2283) was added to the reaction mix at a final concentration of 1 µg/reaction. Unlabeled AbsR28 (35 pmol) and purified *A. baumannii* Hfq72 or *E. coli* Hfq protein (5-fold molar excess in hexamer over *tssM* transcripts) were added to the tubes. $MnCl_2$ was added to the reaction mix at a final concentration of 10 mM. The reaction mixture was incubated at 37 $^{0}$C for 60 min. To initiate the RNase E-mediated degradation, purified RNase E (only the catalytic amino-terminal domain) at 10-fold molar excess over *tssM* transcripts was added to the reaction mixture and incubated further at 37 $^{0}$C for 210 min (0 min denotes the initial time point when RNase E was added). EDTA (2.5 µL from 50 mM stock) and Proteinase K (2.5 µL from 20 mg/mL stock) were added to each reaction mixture and incubated at 50 $^{0}$C for 10 min. Samples were purified immediately using a 2X precipitation buffer supplied with the RNase T1 kit (Ambion, AM2283) according to the manufacturer's instructions. RNA pellets were dissolved in loading buffer II (Ambion, AM2283) and denatured for 3 min at 95° C. RNA cleavage products were resolved on 6% native PAGE at a constant 15 Watt. After gel drying for 2 h at 80 $^{0}$C, bands were visualized using a phosphoimager (Typhoon FLA 9000, GE Healthcare) and ImageQuant software.

**SUPPLEMENTAL FIGURES AND LEGENDS**

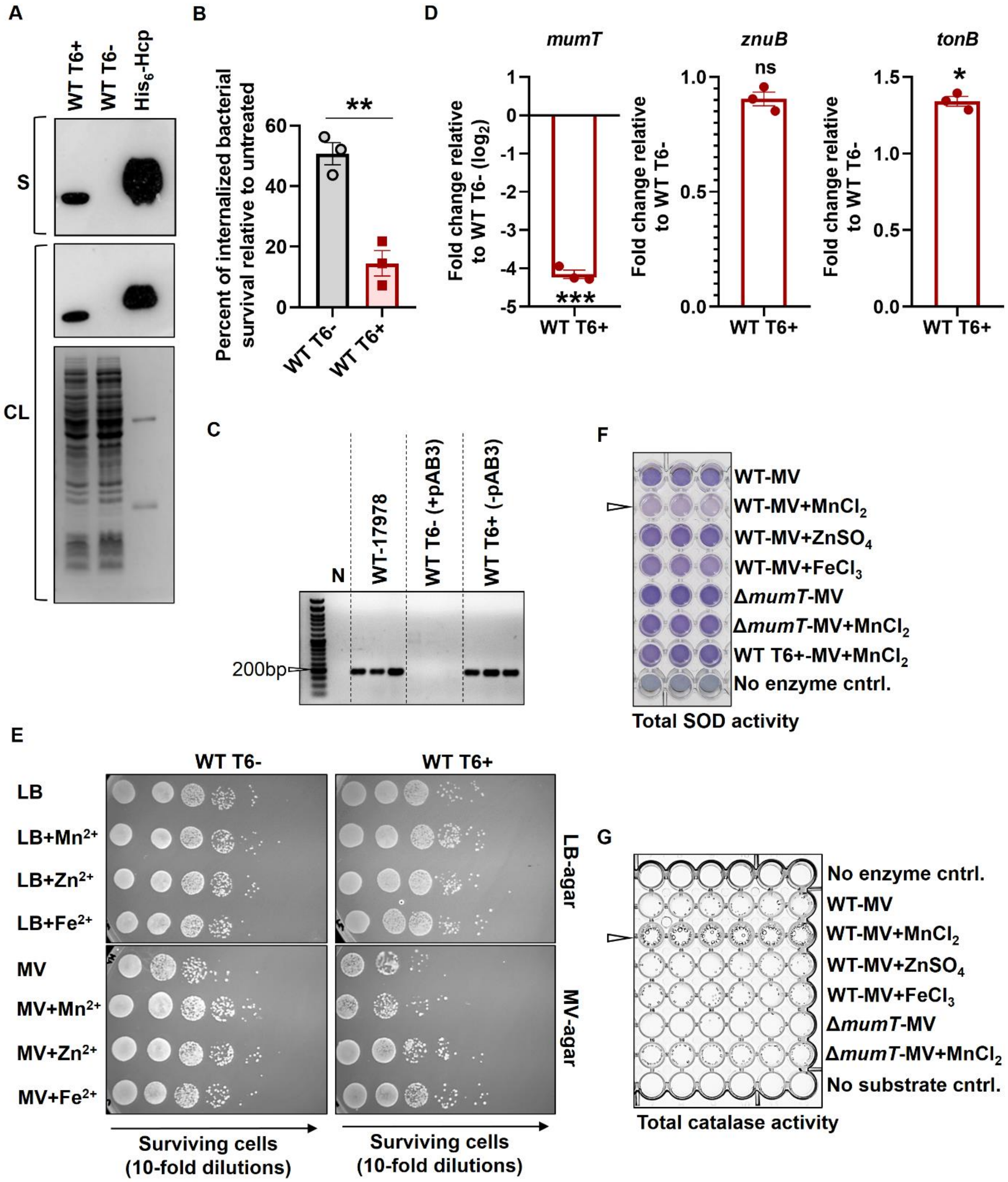


**Figure S1.** *A. baumannii* T6+ cells are sensitive to oxidative stress due to inadequate uptake of $Mn^{2+}$, which is required for SOD and catalase activity. (**A**) The cell-free supernatants (S) of the indicated strains were collected, and the Hcp-secretion profile of the strains was checked by Western blot (upper panel). The Hcp-expression in cell pellet of WT T6- and WT T6+ strains was assessed (lower panel) and gel image below is provided as a loading control. Purified $His_6$-Hcp was used as a positive control for Western Blot. (**B**) Human blood-derived neutrophils were co-incubated with either *A. baumannii* ATCC 17978 wild-type T6SS- (WT T6-) or wild-type T6SS+ (WT T6+) strain for 4 h at an MOI of 1:1. The cells were washed after incubation with gentamycin (300 µg/mL) for 2 h. Neutrophils were lysed using 0.04% Triton X-100. Cell lysates were serially diluted and plated onto Leeds *Acinetobacter* medium plates. The percentage of bacterial survival was enumerated by accounting for the respective untreated control (without phagocytic cells) as 100%. The data represents the mean of biological triplicates ± SEM. Statistical significance was determined using Student's t-test. ** denotes p-value <0.01. (**C**) The presence and absence of *clsc2* (which is present in AbaAL44 island) were checked by PCR in individual colonies of each strain used in the study. Amplification and no amplification for *clsC2* indicate the presence and absence of AbaAL44, respectively. The PCR amplification for *clsC2* was observed in WT T6+ strain whereas no amplification was observed in WT T6- strain. Data representation has been provided for three colonies of each strain. (**D**) The transcript level of *mumT*, *znuB*, and *tonB* was determined by qRT-PCR. The data represents the mean of biological triplicates ± SEM. Statistical significance was determined using Student's t-test. * denotes <0.05, *** denotes p-value <0.001, ns denotes not significant. (**E**) The WT T6- and WT T6+ strains were grown in M9-media containing casamino acid supplemented with either $MnCl_2$, $ZnSO_4$, or $FeCl_3$ (250 µM each) till the mid-log phase ($OD_{600}$~0.6) and incubated further with MV (250 µM) for 2 h. Cells were washed and spotted on LB-agar plates supplemented with MV (100 µM) alone. After incubation at 37 $^0$C overnight (O/N), the images were captured using a gel documentation system (Bio-Rad). (**E**) Total SOD activity was determined using nitroblue tetrazolium (NBT) as a substrate. An equal amount of bacterial cell lysates (grown in M9-media containing casamino acid supplemented with MV + metal ions) were used in this assy. Active SOD causes an inhibition of the photochemical reduction of NBT in the presence of Riboflavin which can be visualized by the naked eye. The highest SOD-mediated inhibition was observed in the presence of $MnCl_2$ (denoted by an empty triangle). (**F**) Total catalase activity was determined using $H_2O_2$ as a substrate. An equal amount of bacterial cell lysates (grown in M9-media containing casamino acid supplemented with MV + metal ions) were used in this assay. Active catalase breaks down $H_2O_2$ to $H_2O$ and $O_2$, which produces air bubbles that can be visualized by the naked eye.

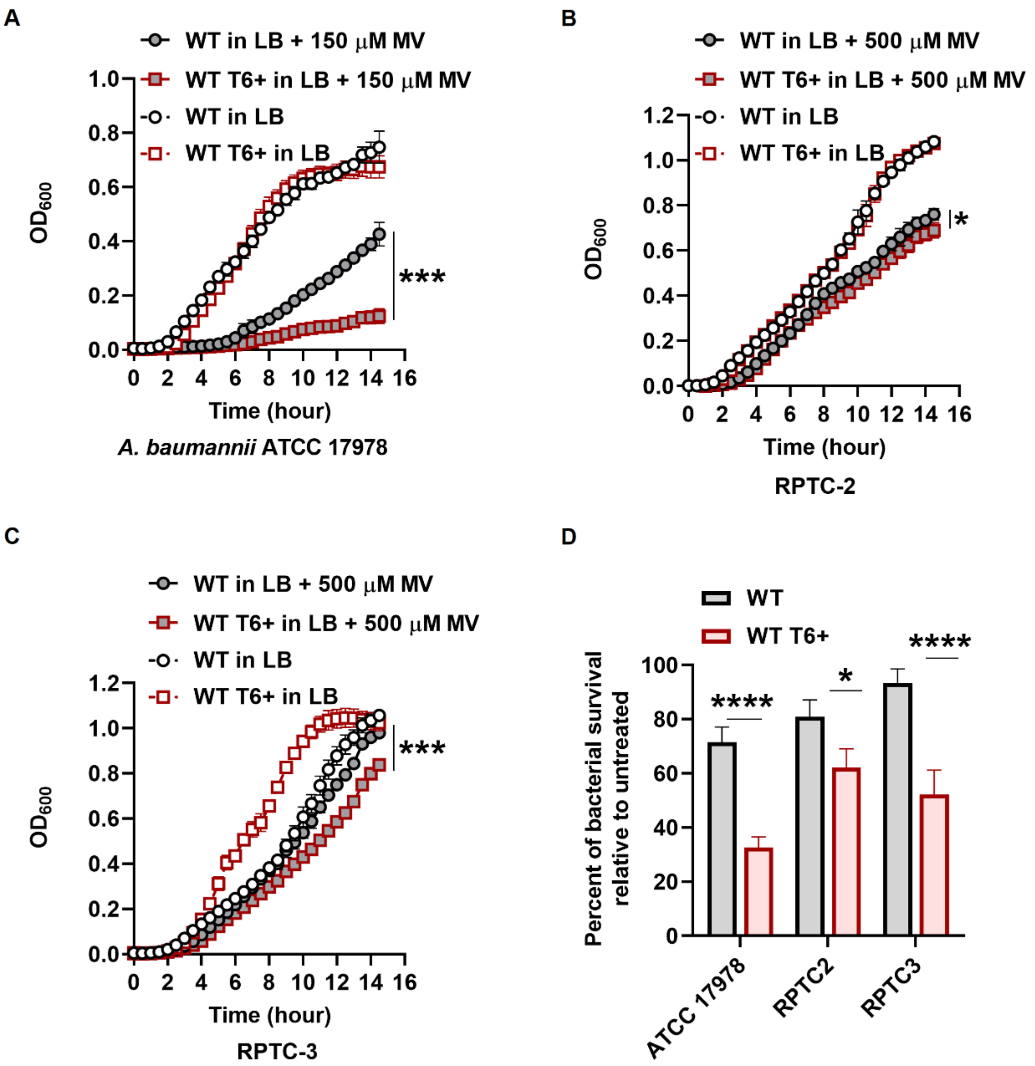


**Figure S2.** The T6+ cells are more susceptible to oxidative stress. (**A-C**) T6+ strains of the bacterial isolates were isolated and compared their growth with their respective wild-type (WT) strain in LB medium supplemented with MV. The data represents mean ± SD. Statistical significance was determined using the Student's t-test. (**D**) Human blood-derived neutrophils were co-incubated with the indicated strains for 4 h at an MOI of 1:1. The percentage of bacterial survival was enumerated by accounting for the respective untreated control (without phagocytic cells) as 100%. The data represents the mean of biological triplicates ± SEM. Statistical significance was determined using the multiple comparison two-way ANOVA test with the Sidak correction for multiple comparisons, comparing the means of each group to one another. * denotes p-value <0.05, **** denotes p-value <0.0001.

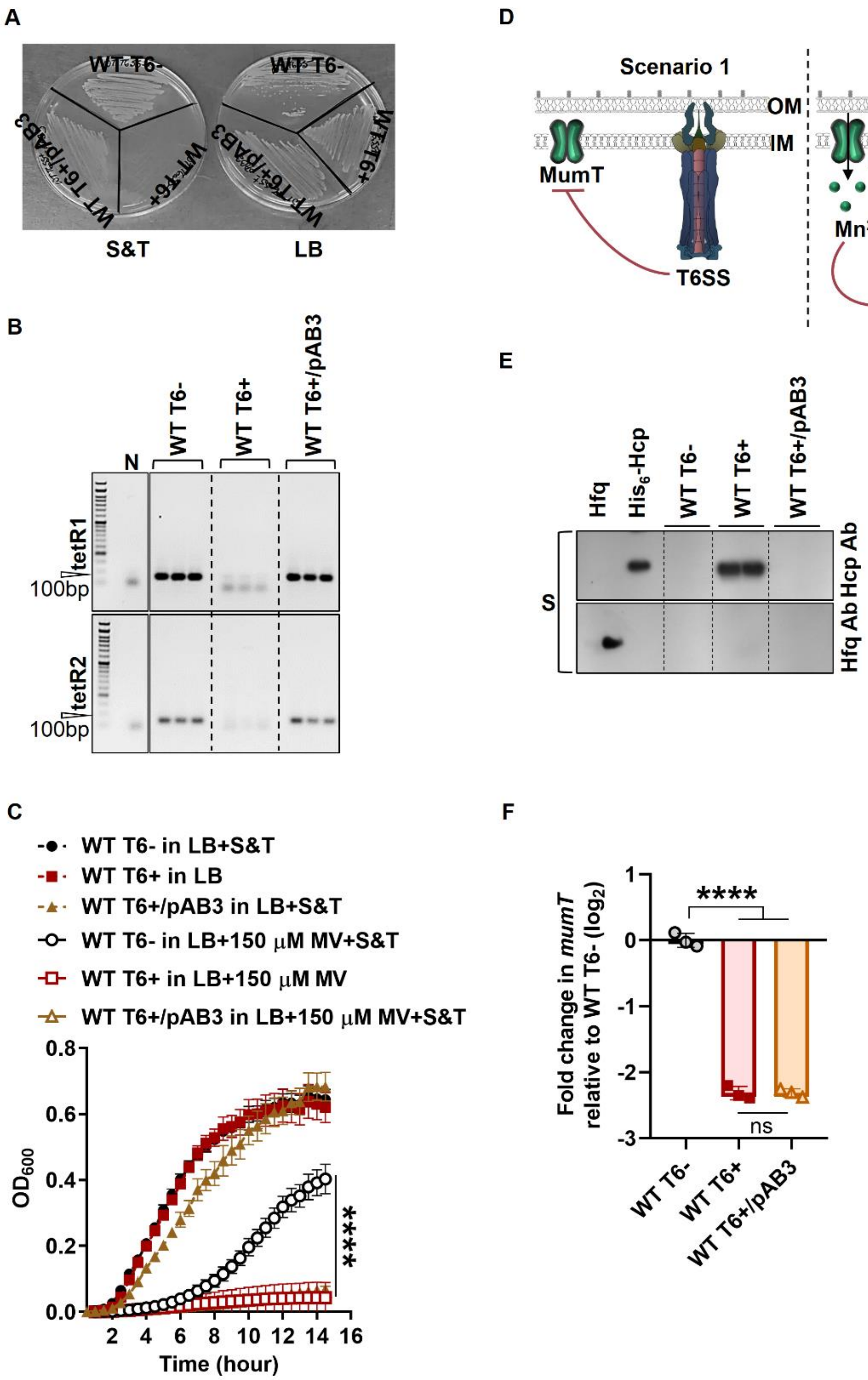


**Figure S3.** pAB3 has no role in mitigating oxidative stress. (**A**) pAB3 was transformed into the WT T6+ competent cells, and the transformants were confirmed by streaking on LB agar plate containing sulfamethoxazole/trimethoprim (S&T; 30 µg/mL and 5 µg/mL, respectively), where the cells lacking pAB3 will not grow. (**B**) The transformation of pAB3 into WT T6+ competent cells was confirmed by PCR using forward and reverse primers of *tetR1* and *tetR2,* which are present in pAB3. Bacterial cells devoid of pAB3 will not show any amplification. (**C**) Growth of WT T6-, WT T6+, and WT T6+/pAB3 cells in LB supplemented with MV (at a final concentration of 150 µM). Except for the WT T6+ cells, ¼ MIC concentration of S/T was added to the medium to maintain the pAB3. Due to addition of S/T, the concentration of MV was kept less. The data represents four biological replicates in technical duplicate with a standard deviation (SD) of the mean. (**D**) Schematic representation of two possible scenarios. (**E**) The cell-free supernatants (S) of the indicated strains were collected, and the Hcp-secretion profile of the strains (in duplicates) was checked by Western blot. Hfq antibody was used to confirm that the supernatants were free from the bacterial cell. (**F**) The transcripts level of *mumT* in WT T6-, WT T6+, and WT T6+/pAB3 strains in LB supplemented with MV was determined by qRT-PCR. The data represents the mean ± SD. Statistical significance was determined using the one-way ANOVA test with Dunnett's multiple comparisons (C) and Tukey's multiple comparisons (E, F). **** denotes p-value <0.0001, ns denotes not significant. ND denotes not detected.

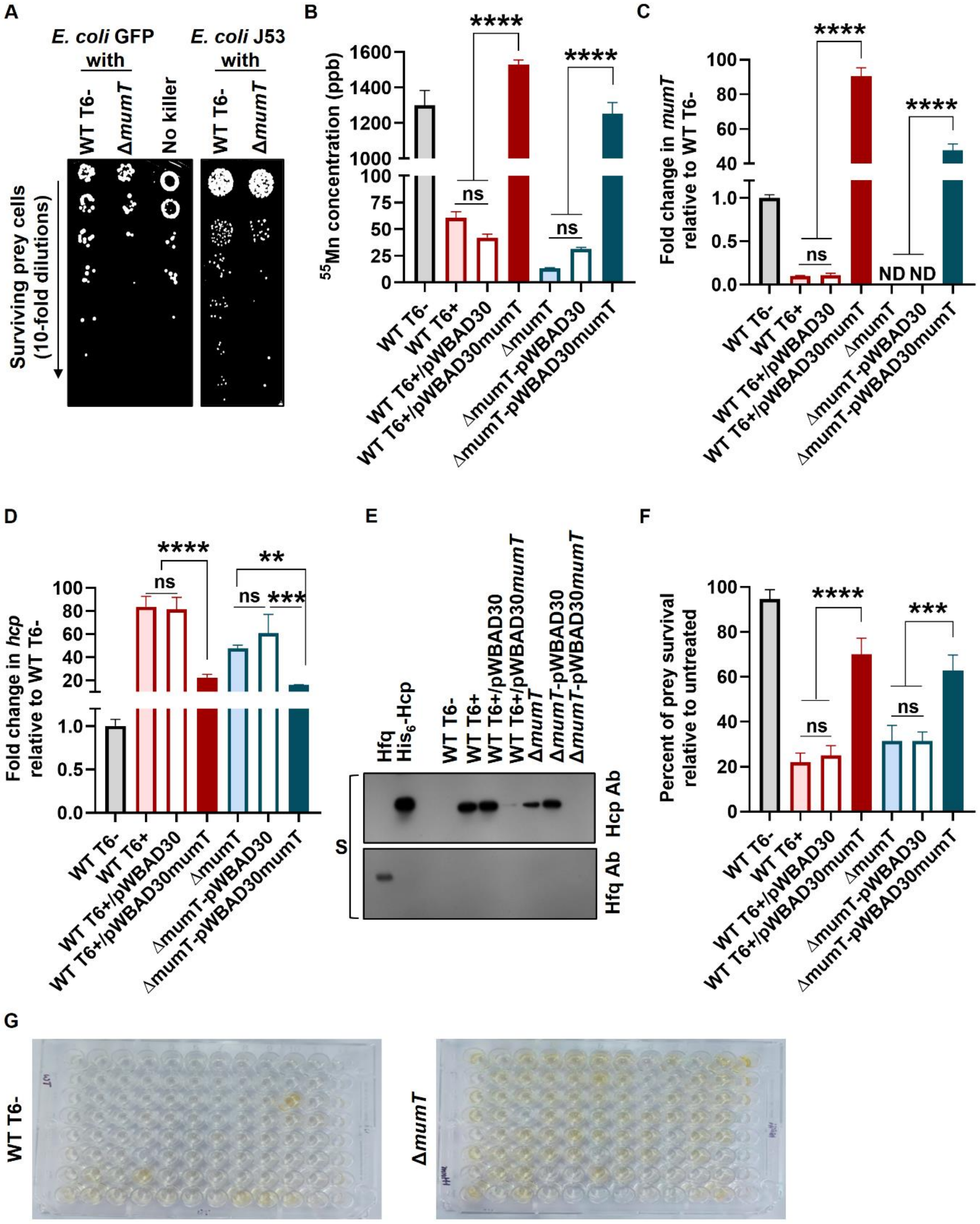


**Figure S4.** Complementation of *mumT* results in a decrease in T6SS expression in *A. baumannii* under oxidative stress. (**A**) Recovery of surviving prey cells after co-incubation with WT T6- or Δ*mumT*. (**B**) Intracellular $^{55}$Mn was quantified by ICP-MS. The bacterial strains were grown in minimal media (M9-media) containing casamino acid as a nutrient source supplemented with MV + $MnCl_2$ (at a final concentration of 100 µM), and the intracellular $Mn^{2+}$ concentration was measured in cell pellets by ICP-MS. The data represents three biological replicates with a standard deviation (SD) of the mean. (**C & D**) The bacterial strains were grown in LB supplemented with $MnCl_2$ (250 µM) till the mid-log phase ($OD_{600}$~0.6) and incubated further with MV (250 µM). RNA was extracted from the cells after arabinose induction, and cDNA was prepared. The expression level of *hcp* and *mumT* transcripts in the strains was determined by qRT-PCR. The data represents the mean ± SD. (**E**) The cell-free supernatants (S) from the above mentioned experiment were collected, and the Hcp-secretion profile of the strains was checked by Western blot. Hfq antibody was used to confirm that the supernatants were free from the bacterial cell. (**F**) In the T6SS competition assay, prey cells (*E.coli* J53) were subjected to killing by incubation with the indicated strains. The survival percentage of prey cells was enumerated by accounting for the respective untreated control (without predator/killer cells) as 100%. The data represents the mean of three biological replicates ± SD. For all the assays, WT T6- and WT T6+ cells were used as T6- and T6+ control, respectively. Statistical significance was determined using the one-way ANOVA test with Tukey's multiple comparisons. ** denotes p-value <0.01, *** denotes p-value <0.001, **** denotes p-value <0.0001, ns denotes not significant. ND denotes not detected. (**G**) Original pictures are captured by a camera that is schematically represented in Figure 2E.

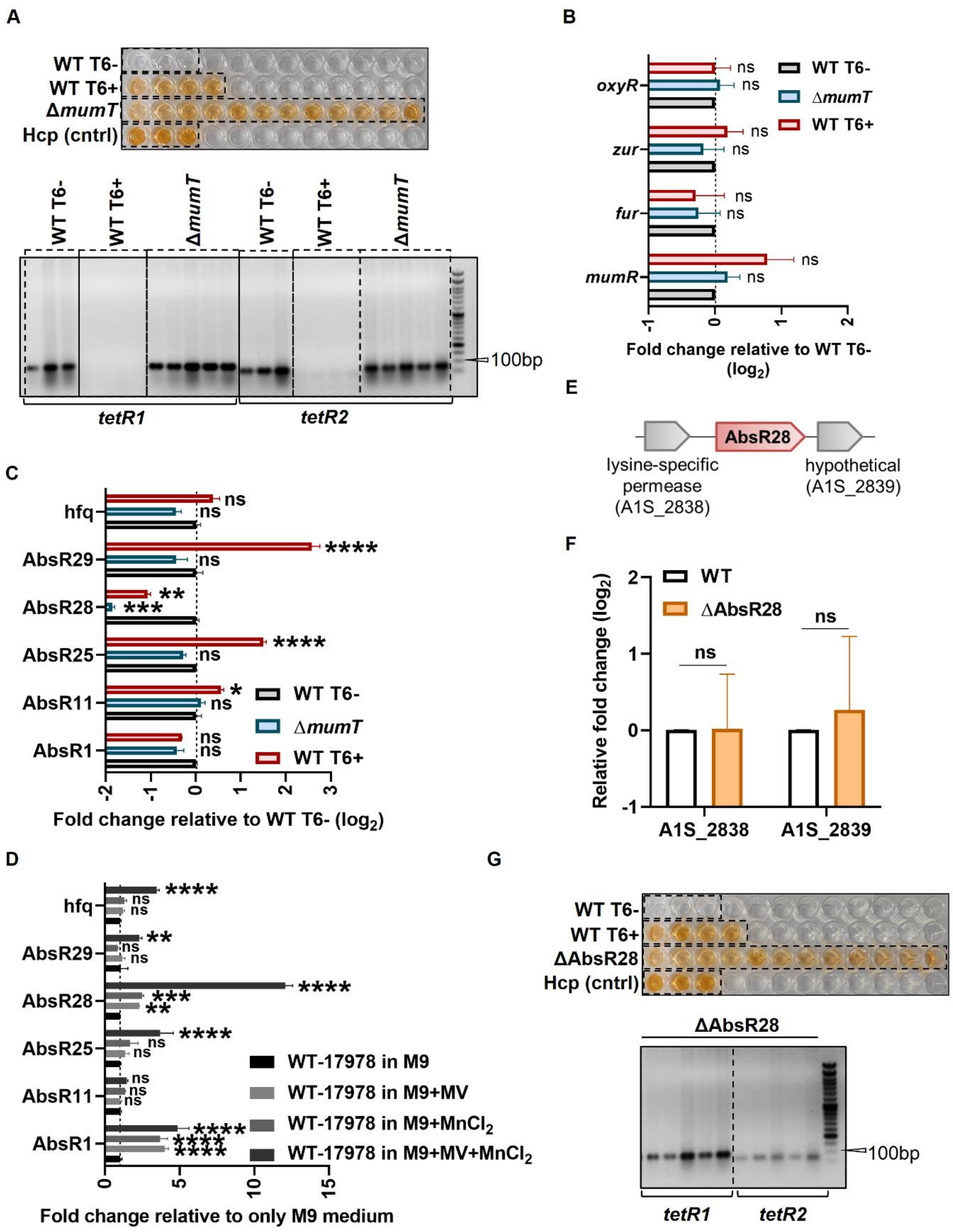


**Figure S5.** Supplementation of $MnCl_2$ results in an increase of AbsR28 at the transcript level. (**A**) Hcp-ELISA assay was performed for Δ*mumT* strain (upper panel) and wells that exhibited Hcp-secretion positive were picked up to determine the presence of pAB3 by PCR using forward and reverse primers of *tetR1* and *tetR2*, which are present in pAB3. Bacterial cells devoid of pAB3 will not show any amplification. WT T6- and WT T6+ strains were used as pAB3 positive and pAB3 negative control for this assay, respectively. Gel image of five samples is represented. (**B**) The expression of several transcriptional regulators of T6SS was determined by qRT-PCR, where the bacterial strains were grown in an LB medium. The data represents the mean ± SD. (**C**) The expression of several sRNAs and *hfq* was determined by qRT-PCR, where the bacterial strains were grown in an LB medium. The data represents the mean ± SD. (**D**) The expression of several sRNAs and *hfq* was determined by qRT-PCR, where the bacterial strains were grown in M9-media containing casamino acid supplemented with either MV or $MnCl_2$ alone or MV+$MnCl_2$ (250 µM). (**E**) Genomic location of AbsR28 in *A. baumannii* ATCC 17978 (NCBI Ref. seq. CP000521.1). (**F**) To test the polar effect, WT and ΔAbsR28 strains were grown in LB supplemented with $MnCl_2$ (250 µM) till the mid-log phase ($OD_{600}$~0.6) and incubated further with MV (250 µM) for 2 h. RNA was extracted from the cells, and cDNA was prepared. The expression of A1S_2828 and A1S_2839 transcripts, which are immediate upstream and downstream of AbsR28, respectively, was checked by qRT-PCR. No change in the expression in ΔAbsR28 for both the genes with respect to WT confirms no polar effect of ΔAbsR28. (**G**) Hcp-ELISA assay was performed for ΔAbsR28 strain (upper panel) and wells that exhibited Hcp-secretion positive were picked up to determine the presence of pAB3 by PCR using forward and reverse primers of *tetR1* and *tetR2*, which are present in pAB3. Bacterial cells devoid of pAB3 will not show any amplification. Gel image of five samples is represented. The data represents the mean ± SD. Statistical significance was determined using the multiple comparison two-way ANOVA test with the Sidak correction for multiple comparisons, comparing the means of each group to one another (C, D). * denotes p-value <0.05, ** denotes p-value <0.01, *** denotes p-value <0.001, **** denotes p-value <0.0001, ns denotes not significant.

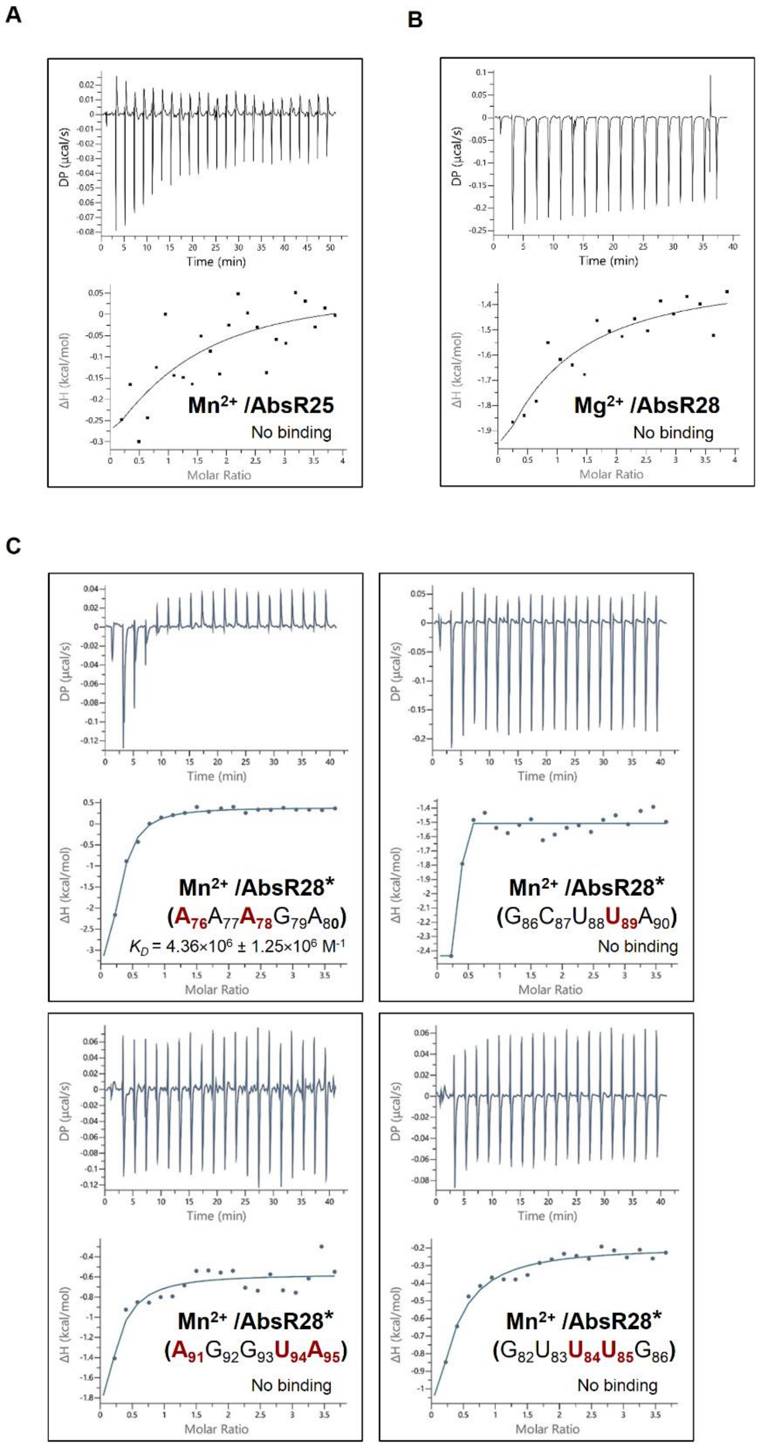


**Figure S6. $Mn^{2+}$ binds to AbsR28.** (**A**) Isothermal titration calorimetry (ITC) of $Mn^{2+}$ to AbsR25 as nonspecific sRNA control. (**B**) ITC of $Mg^{2+}$ to AbsR28 as nonspecific metal ion control. (**C**) ITC of $Mn^{2+}$ to AbsR28 mutants. Mutated nucleotides are highlighted.

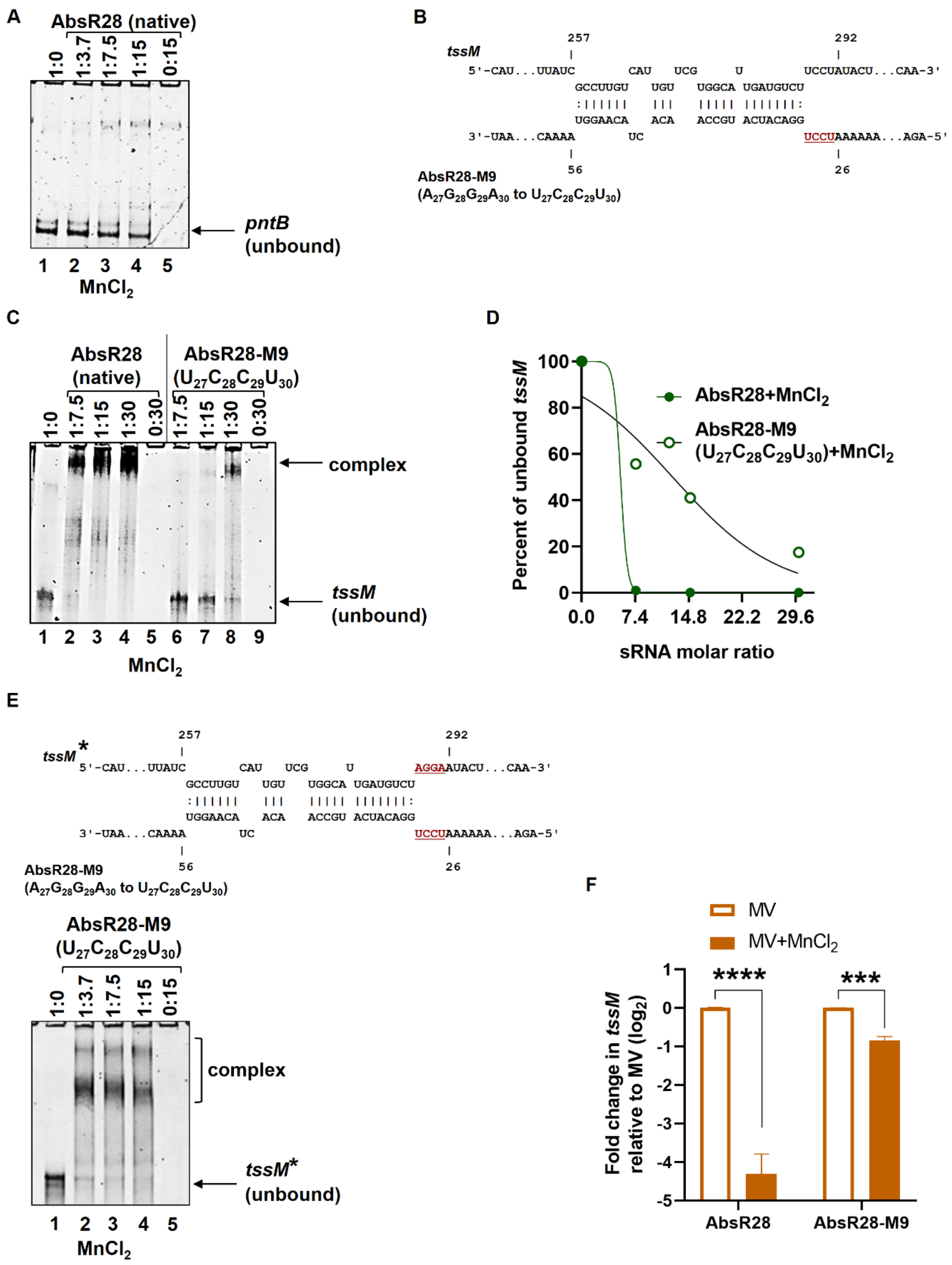


**Figure S7.** AbsR28 base pairs with *tssM* mRNA in the presence of $Mn^{2+}$. (**A**) Gel retardation assay of unlabeled *pntB in vitro* transcripts (used as a negative mRNA control)

with unlabeled full-length AbsR28 *in vitro* transcripts in structure buffer containing $MnCl_2$. (**B**) The position of the four nucleotides of the seed region of AbsR28 is mutated (AbsR28-M9) and shown in red underlined. (**C**) Gel retardation assay of unlabeled *tssM in vitro* transcripts with unlabeled full-length AbsR28 or AbsR28-M9 *in vitro* transcripts in structure buffer containing $MnCl_2$. (**D**) Quantification of unbound *tssM* obtained from Figure S6C (n = 2 independent experiments) is shown using ImageJ software. Nonlinear regression was used to fit the curve. (**E**) Gel retardation assay of unlabeled compensatory mutation in *tssM* mRNA (*tssM**) *in vitro* transcripts with unlabeled AbsR28-M9 *in vitro* transcripts in structure buffer containing $MnCl_2$. (**F**) The relative level of *tssM* transcripts under the arabinose pulse was determined by qRT-PCR. The data represents the mean ± SD. Statistical significance was determined using the multiple comparison two-way ANOVA test with the Sidak correction for multiple comparisons, comparing the means of each group to one another. *** denotes p-value <0.001, **** denotes p-value <0.0001.

**Table S1.** CopraRNA search results for each of the 31 sRNAs (whose sequences were bioinformatically predicted) that showed T6SS genes as hit.

| small RNA | Component of T6SS | Binding energy |
|---|---|---|
| AbsR1 | type VI secretion system ATPase TssH | -7.49 |
| AbsR2 | Nil | NA |
| AbsR3 | Nil | NA |
| AbsR4 | type VI secretion system baseplate subunit TssG | -9.47 |
| AbsR5 | type VI secretion system protein TssA | -15.12 |
| AbsR6 | Nil | NA |
| AbsR7 | Nil | NA |
| AbsR8 | type VI secretion system protein TssA<br>type VI secretion system baseplate subunit TssG | -8.33<br>-7.58 |
| AbsR9 | Nil | NA |
| AbsR10 | Nil | NA |
| AbsR11 | type VI secretion system-associated protein TagF | -14.58 |
| AbsR12 | type VI secretion system baseplate subunit TssK<br>type VI secretion system-associated protein TagF<br>type VI secretion system tube protein Hcp<br>type VI secretion system baseplate subunit TssG | -11.06<br>-8.92<br>-8.72<br>-8.71 |
| AbsR13 | type VI secretion system baseplate subunit TssK | -15.64 |
| AbsR14 | type VI secretion system contractile sheath large subunit TssC | -22.86 |
| AbsR15 | type VI secretion protein TssM | -14.76 |
| AbsR16 | Nil | NA |
| AbsR17 | type VI secretion system-associated protein TagF | -12.23 |
| AbsR18 | type VI secretion protein TssM | -9.37 |
| AbsR19 | Nil | NA |
| AbsR20 | type VI secretion system protein | -13.31 |
| AbsR21 | type VI secretion system baseplate subunit TssG<br>type VI secretion system protein | -14.7<br>-10.76 |
| AbsR22 | Nil | NA |
| AbsR23 | Nil | NA |
| AbsR24 | Nil | NA |
| AbsR25 | type VI secretion system baseplate subunit TssK<br>type VI secretion system protein TssA | -14.15<br>-13.57 |
| AbsR26 | type VI secretion system baseplate subunit TssK | -9.44 |
| AbsR27 | Nil | NA |
| AbsR28 | type VI secretion protein TssM<br>type VI secretion system baseplate subunit TssG | -18.29<br>-10.54 |
| AbsR29 | type VI secretion system protein<br>type VI secretion system-associated protein TagF | -14.9<br>-12.73 |
| AbsR30 | Nil | NA |
| AbsR31 | type VI secretion system baseplate subunit TssK | -15.64 |

**Table S2.** Genes with altered expression ($log_2$<-1 and >+1) in ΔAbsR28 strain relative to WT T6- grown in LB medium supplemented with MV+$MnCl_2$, both at final conc. of 250 µM (represented in Figure 3B).

| Gene locus | Annotation (KEGG) and function | Fold change ($log_2$) |
|---|---|---|
| A1S_2668 | Phosphoenolpyruvate carboxykinase (pyruvate metabolism) | -1.145 |
| A1S_0804 | Trehalose-6-phosphate phosphatase (Starch and sucrose metabolism) | 1.718 |
| A1S_0140 | NAD-linked malate dehydrogenase (pyruvate metabolism) | 1.583 |
| A1S_1341 | Enoyl-CoA hydratase/carnithine racemase (fatty acid degradation) | 1.026 |
| A1S_3362 | Hypothetical protein (glycerophospholipid metabolism) | 2.034 |
| A1S_0041 | Putative linoleoyl-CoA desaturase (biosynthesis of unsaturated fatty acids) | 1.279 |
| A1S_3356 | Putative flavoprotein monooxygenase (purine metabolism) | -1.326 |
| A1S_2686 | Carbamoyl-phosphate synthase small chain (pyrimidine metabolism) | -1.033 |
| A1S_1528 | Hypothetical protein (amino acid metabolism) | -1.152 |
| A1S_0971 | Methionine synthase (cysteine and methionine metabolism) | -3.113 |
| A1S_0737 | 5-Methyltetrahydropteroyltriglutamate-homocysteine methyltransferase (cysteine and methionine metabolism) | -1.213 |
| A1S_1376 | Acyl-CoA dehydrogenase (amino acid degradation) | -1.238 |
| A1S_1341 | Enoyl-CoA hydratase/carnithine racemase (amino acid degradation) | 1.026 |
| A1S_1375 | Putative propionyl-CoA carboxylase (β subunit) (amino acid degradation) | -1.082 |
| A1S_1277 | Allophanate hydrolase subunit 2 (arginine biosynthesis) | -1.312 |
| A1S_1283 | Putative amidase (arginine biosynthesis) | -1.543 |
| A1S_1528 | Hypothetical protein (arginine and proline metabolism) | -1.152 |
| A1S_3415 | Maleylacetoacetate isomerase (tyrosine metabolism) | 1.121 |
| A1S_3414 | Fumarylacetoacetase (tyrosine metabolism) | 1.015 |
| A1S_1336 | Hypothetical protein (phenylalanine metabolism) | 1.758 |
| A1S_1337 | Phenylacetic acid degradation B (phenylalanine metabolism) | 1.61 |
| A1S_1338 | Hypothetical protein (phenylalanine metabolism) | 1.832 |
| A1S_1340 | Phenylacetate-CoA oxygenase/reductase PaaK subunit (phenylalanine metabolism) | 1.26 |
| A1S_1335 | Phenylacetic acid degradation protein paaN (phenylalanine metabolism) | 1.138 |
| A1S_1341 | Enoyl-CoA hydratase/carnithine racemase (phenylalanine metabolism) | 1.026 |
| A1S_1075 | D-amino-acid dehydrogenase (phenylalanine metabolism) | 1.268 |
| A1S_1341 | Enoyl-CoA hydratase/carnithine racemase (phenylalanine metabolism) | 1.026 |
| A1S_2448 | Putative phosphate transporter (two-component system) | 1.628 |
| A1S_2301 | Amino acid ABC transporter permease protein (ABC transporter) | -1.425 |
| A1S_0466 | Sec-independent protein translocase protein (twin-arginine translocation system) | 1.205 |
| A1S_2448 | Putative phosphate transporter (two-component system) | 1.628 |
| A1S_0140 | NAD-linked malate dehydrogenase (two-component system) | 1.583 |
| A1S_1978 | Response regulator protein (two-component system) | -1.021 |
| A1S_2196 | Membrane-associated dicarboxylate transport protein (two-component system) | 1.344 |
| A1S_1925 | Cytochrome d terminal oxidase polypeptide subunit II (two-component system) | -1.073 |
| A1S_1384 | CinA-like protein (nicotinate and nicotinamide metabolism) | 1.627 |
| A1S_0566 | Pyridine nucleotide transhydrogenase (proton pump) α subunit (part1) (nicotinate and nicotinamide metabolism) | -2.918 |

| | | |
|---|---|---|
| A1S_0567 | Pyridine nucleotide transhydrogenase (proton pump) α subunit (part2) (nicotinate and nicotinamide metabolism) | -3.028 |
| A1S_0568 | Pyridine nucleotide transhydrogenase β subunit (nicotinate and nicotinamide metabolism) | -2.95 |
| A1S_0466 | Sec-independent protein translocase protein (protein export) | 1.205 |
| A1S_1221 | Hypothetical protein (mismatch repair) | 1.118 |

**Table S3.** Bacterial strains were used in this study.

| **Strains** | **Name used in this study** | **Description** | **Reference** |
|---|---|---|---|
| *Acinetobacter baumannii* ATCC 17978 | WT | Wild-type (WT) strain, AbaAL44+ | ATCC, USA |
| *Acinetobacter baumannii* ATCC 17978 T6- | WT T6- | T6SS repressed, pAB3 present, $S/T^R$ | This study |
| *Acinetobacter baumannii* ATCC 17978 T6+ | WT T6+ | T6SS expressed, pAB3 absent | This study |
| *Acinetobacter baumannii* ATCC 17978 T6+/ pWBAD30 | WT T6+/ pWBAD30 | WT T6+ cells with pWBAD30, pAB3 absent, $Kan^R$ | This study |
| *Acinetobacter baumannii* ATCC 17978 T6+/ pWBAD30*mumT* | WT T6+/ pWBAD30*mumT* | WT T6+ cells with pWBAD30*mumT*, pAB3 absent, $Kan^R$ | This study |
| *Acinetobacter baumannii* ATCC 17978 Δ*mumT* | Δ*mumT* | *mumT* k/o in *A. baumannii* ATCC 17978, pAB3 present | This study |
| *Acinetobacter baumannii* ATCC 17978 Δ*mumT*- pWBAD30 | Δ*mumT*- pWBAD30 | *mumT* k/o in *A. baumannii* ATCC 17978, pAB3 present, $Kan^R$ | This study |
| *Acinetobacter baumannii* ATCC 17978 Δ*mumT*- pWBAD30mumT | Δ*mumT*- pWBAD30mumT | *mumT* k/o in *A. baumannii* ATCC 17978, pAB3 present, $Kan^R$ | This study |
| *Acinetobacter baumannii* ATCC 17978 ΔAbsR28 | ΔAbsR28 | ΔAbsR28 k/o in *A. baumannii* ATCC 17978, pAB3 present | This study |
| *Acinetobacter baumannii* ATCC 17978 ΔAbsR28-pWBAD30 | ΔAbsR28- pWBAD30 | ΔAbsR28 k/o in *A. baumannii* ATCC 17978 with pWBAD30, pAB3 present, $Kan^R$ | This study |
| *Acinetobacter baumannii* ATCC 17978 ΔAbsR28-pWBAD30AbsR28 | ΔAbsR28- pWBAD30AbsR28 | ΔAbsR28 k/o in *A. baumannii* ATCC 17978 with pWBAD30AbsR28, pAB3 present, $Kan^R$ | This study |
| *Acinetobacter baumannii* ATCC 17978 ΔAbsR28-pWBAD30AbsR28 mutant_M9 ($U_{27}C_{28}C_{29}U_{30}$) | ΔAbsR28- pWBAD30AbsR28 mutant_M9 ($U_{27}C_{28}C_{29}U_{30}$) | ΔAbsR28 k/o in *A. baumannii* ATCC 17978 with pWBAD30AbsR28 mutant_M9 ($U_{27}C_{28}C_{29}U_{30}$), pAB3 present, $Kan^R$ | This study |
| *Acinetobacter baumannii* ATCC 17978 Δ*tssM* | Δ*tssM* | *tssM* k/o in *A. baumannii* ATCC 17978, pAB3 present | Lab stock |
| *Acinetobacter baumannii* ATCC 17978 T6+/pAB3 | WT T6+/pAB3 | T6SS repressed, pAB3 present, $S/T^R$ | This study |
| *Acinetobacter baumannii* RPTC2 | RPTC2 | Clinical isolate | Dr. Varsha Gupta, GMCH, |

| | | | |
|---|---|---|---|
| | | | Chandigarh, India |
| *Acinetobacter baumannii* RPTC2 T6+ | RPTC2 T6+ | T6SS expressed | This study |
| *Acinetobacter baumannii* RPTC3 | RPTC3 | Clinical isolate | Dr. Varsha Gupta, GMCH, Chandigarh, India |
| *Acinetobacter baumannii* RPTC3 T6+ | RPTC3 T6+ | T6SS expressed | This study |
| *Escherichia coli* DH5α | *E. coli* DH5α | *supE44 hsdR17 recA1 endA1 gyrA96 thi-1 relA1* | Invitrogen, USA |
| *Escherichia coli* DH5α-pNYL GFP | *E. coli*-pNYL GFP | *E. coli*-pNYL GFP | Prof. N.K. Navani, IIT Roorkee, India |
| *Escherichia coli* J53 | *E. coli* J53 | *E. coli* J53 | Dr. Sanath Kumar H, ICAR-CIFE, India |
| *Pseudomonas aeruginosa* PAO1 | *P. aeruginosa* | *P. aeruginosa* | Prof. N.K. Navani, IIT Roorkee, India |

**Table S4.** Plasmids were used in this study.

| **Strains** | **Name used in this study** | **Description** | **Reference** |
|---|---|---|---|
| pUC18 | N/A | Cloning vector, $Amp^R$ | Thermo Scientific, USA |
| pMDIAI | N/A | Plasmid carrying apramycin (Apm) resistance cassette flanked by FRT sites | Addgene |
| pUC18-UP*mumT*-AprFRT-DN*mumT* | N/A | *mumT* construct for *mumt* k/o cloned in pUC18, $Amp^R$, $Apm^R$ | This study |
| pUC18-UPAbsR28-AprFRT-DNAbsR28 | N/A | AbsR28 construct for AbsR28 k/o cloned in pUC18, $Amp^R$, $Apm^R$ | This study |
| pAT02 | N/A | Plasmid expressing *A. baumannii* RecT homolog, $Amp^R$ | Prof. Bryan Davies, University of Texas, San Antonio, USA |
| pAT03 | N/A | Plasmid expressing FLP recombinase enzyme (flippase) for expression in *A. baumannii*, $Amp^R$ | Prof. Bryan Davies, University of Texas, San Antonio, USA |
| pBAD30-$Amp^R$ | N/A | Arabinose PBAD promoter, $Amp^R$ | Prof. Eric D. Brown, McMaster University, Hamilton, ON |

| | | | |
|---|---|---|---|
| pBAD30-Kan$^R$ | pBAD30 | Modified from pBAD30-Amp$^R$, arabinose PBAD promoter, Kan$^R$ | This study |
| pWBAD30-Kan$^R$ | pWBAD30 | Modified from pBAD30-Kan$^R$, arabinose PBAD promoter, A. baumannii compatible *ori* cloned, Kan$^R$ | This study |
| pWBAD30-Kan$^R$-AbsR28 | pWBAD30-AbsR28 | AbsR28 cloned under PBAD promoter, Kan$^R$ | This study |
| pWBAD30-Kan$^R$-mumT | pWBAD30-mumT | *mumT* cloned under PBAD promoter, Kan$^R$ | This study |
| pWBAD30-Kan$^R$-AbsR28 mutant-M9 ($U_{27}C_{28}C_{29}U_{30}$) | pWBAD30AbsR28 mutant-M9 ($U_{27}C_{28}C_{29}U_{30}$) | AbsR28 mutant-M9 cloned under PBAD promoter, Kan$^R$ | This study |
| pUC18-M12 | N/A | AbsR28 mutant ($C_{76}C_{78}$ to $A_{76}A_{78}$) cloned in pUC18 | This study |
| pUC18-M14 | N/A | AbsR28 mutant ($C_{89}$ to $U_{89}$) cloned in pUC18 | This study |
| pUC18-M15 | N/A | AbsR28 mutant ($C_{91}A_{94}C_{95}$ to $A_{91}U_{94}A_{95}$) cloned in pUC18 | This study |
| pUC18-M20 | N/A | AbsR28 mutant ($C_{84}A_{85}$ to $U_{84}U_{85}$) cloned in pUC18 | This study |
| pUC18-*tssM** | N/A | Compensatory *tssM* mutant ($C_{84}A_{85}$ to $U_{84}U_{85}$) cloned in pUC18 | This study |

**Table S5.** Oligonucleotides were used in this study.

| **Primer name** | **Sequence (5' - 3')** | **Description** | **Reference** |
|---|---|---|---|
| UP437bpFPmumT | ATGCGTCGACAATTAACTGAAGTGGC | Forward primer for cloning *mumT* 437 bp upstream into pUC18 | This study |
| UP437bpRPmumT | ACTCTAGATTAATGCGTTCCTCATCCATTTG | Reverse primer for cloning *mumT* 437 bp upstream into pUC18 | This study |
| DN500bpFPmumT | ATGGTACCAGACGACATGAATTGATAAG | Forward primer for cloning *mumT* 500 bp downstream into pUC18 | This study |
| DN500bpRPmumT | ATGGAATTCAACTCGTGCTTGCTCG | Reverse primer for cloning *mumT* 500 bp downstream into pUC18 | This study |
| UP125bpFPmumT | ATGGGGAACAGGGAATCTTTGTCAT | Forward primer for the amplification of *mumT* 125 bp upstream | This study |
| DN125bpRPmumT | ACCAGCATGAAAACCACAAGCAAT | Reverse primer for the amplification of *mumT* 125 bp downstream | This study |
| UP498bpFPAbsR28 | TAAGTCGACATATGCAACTACATTCATTGCTGC | Forward primer for cloning AbsR28 498 bp upstream into pUC18 | This study |
| UP498bpRPAbsR28 | TTATCTAGAGAACGGATTTTACCTGTTTT | Reverse primer for cloning AbsR28 498 bp upstream into pUC18 | This study |
| DN501bpFPAbsR28 | TATGGTACCAAATAAGAGAATAATTATGGGCAT | Forward primer for cloning AbsR28 501 bp downstream into pUC18 | This study |
| DN501bpRPAbsR28 | TATGAATTCCCTAAAGTGCCCAGCTGTTTT | Reverse primer for cloning AbsR28 501 bp downstream into pUC18 | This study |

| | | | |
|---|---|---|---|
| UP126bpFPAbsR28 | ATTAATTCCTTACGATCAAATGGATGTAAAACC | Forward primer for the amplification of AbsR28 126 bp upstream | This study |
| DN126bpRPAbsR28 | TGTGCCATTTTCTTGAGTTGTTCAATACTT | Reverse primer for the amplification of AbsR28 126 bp downstream | This study |
| AprF-Bam | ATCAGGATCCGTCGACCTGCAGTTC | Forward primer for cloning Apr-FRT into pUC18 | Lab stock |
| AprR-Kpn | ATGGTACCGTGTAGGCTGGAGCTGCTTC | Reverse primer for cloning Apr-FRT into pUC18 | Lab stock |
| Kan FP pWBAD30 | TTCGATCGGAACTTCAAGATCCCCTCAC | Forward primer for cloning Kanamycin resistance marker into pBAD30 | Lab stock |
| Kan RP pWBAD30 | TTCGATCGTTCTCGAGAAGTATAGGAACTTCAGAGC | Reverse primer for cloning Kanamycin resistance marker containing XhoI restriction site into pBAD30 | Lab stock |
| pW FP | TGTCTCGAGGATCGTAGAAATATCTATGATTATC | Forward primer for cloning *A. baumannii* ori into pBAD30-kan$^R$ | Lab stock |
| pW RP | TGTCTCGAGGGATTTTAACATTTTGCGTTGTTC | Reverse primer for cloning *A. baumannii* ori into pBAD30-kan$^R$ | Lab stock |
| AbsR28 FP pWBAD30 | AGGAATTCGTCAAAAACTTGATCTTTAG | Forward primer for cloning AbsR28 into pWBAD30-kan$^R$ | This study |
| AbsR28 RP pWBAD30 | CCCAAGCTTATTGTCCGAATAGGAATAAAAAAACCTAGCG | Reverse primer for cloning AbsR28 into pWBAD30-kan$^R$ | This study |
| RT *mumT* FP | TGCCATGGATAAAAGCATGA | qRT-PCR forward primer for *mumT* | This study |
| RT *mumT* RP | CAACTGTACCGCCAACAATGG | qRT-PCR reverse primer for *mumT* | This study |
| RT *znuB* FP | ATTTGAGGCTGCCAATAGCG | qRT-PCR forward primer for *znuB* | This study |
| RT *znuB* RP | AAACGATGCTTTGCTTGCCC | qRT-PCR reverse primer for *znuB* | This study |
| RT *tonB* FP | CCAGATCCATCGCCAAAACG | qRT-PCR forward primer for *tonB* | This study |
| RT *tonB* RP | GGGTTACGCGCACGTTAGTA | qRT-PCR reverse primer for *tonB* | This study |
| RT *tssB* FP | TCAGCGAATTCGACCTCCAC | qRT-PCR forward primer for *tssB* | Lab stock |
| RT *tssB* RP | GTACGCTCAAGCTCAGATGC | qRT-PCR reverse primer for *tssB* | Lab stock |
| RT *tssC* FP | GTTGGTGTGCTGCTATTCGC | qRT-PCR forward primer for *tssC* | This study |
| RT *tssC* RP | CTCTTTTTCACGGCGATCCG | qRT-PCR reverse primer for *tssC* | This study |
| RT *hcp* FP | CTTCAAGTAGTGTAGGCGGC | qRT-PCR forward primer for *hcp* | Lab stock |
| RT *hcp* RP | CCATTTGCACGATAGAAGTC | qRT-PCR reverse primer for *hcp* | Lab stock |
| RT *tssE* FP | GTGGGGCTTTCTACAGCCAA | qRT-PCR forward primer for *tssE* | This study |
| RT *tssE* RP | ACCCGTATTTGTCTTAGCCGAG | qRT-PCR reverse primer for *tssE* | This study |

| | | | |
|---|---|---|---|
| RT *tssF* FP | TAGTAGCTTGGCGA GACGTG | qRT-PCR forward primer for *tssF* | Lab stock |
| RT *tssF* RP | GATCACACGCCACT GTTCAC | qRT-PCR reverse primer for *tssF* | Lab stock |
| RT *tssG* FP | ACCTGGTGCAGTCCA ACTTT | qRT-PCR forward primer for *tssG* | This study |
| RT *tssG* RP | AAAAAGCGCCTTGC CCTAAG | qRT-PCR reverse primer for *tssG* | This study |
| RT *tssM* FP | CTCCGGCAACCAATC AGTCT | qRT-PCR forward primer for *tssM* | This study |
| RT *tssM* RP | AGCTGTAATACGAG CACCCG | qRT-PCR reverse primer for *tssM* | This study |
| RT *paar* FP | TGGCTAGCCCTTACA TTACG | qRT-PCR forward primer for *paar* | Lab stock |
| RT *paar* RP | CGTTTTATGCGCCGG ACAAG | qRT-PCR reverse primer for *paar* | Lab stock |
| RT *tssH* FP | CTCGAGTGCAATTAT GCAGGC | qRT-PCR forward primer for *tssH* | This study |
| RT *tssH* RP | CACAACTCTCATGCG CCCTA | qRT-PCR reverse primer for *tssH* | This study |
| RT *tssA* FP | CAATCGCGAGCAAG CAATGA | qRT-PCR forward primer for *tssA* | This study |
| RT *tssA* RP | GCTAACCATTCATGC AGCGG | qRT-PCR reverse primer for *tssA* | This study |
| RT *tssK* FP | GCAGACCCACGAGT TGATTC | qRT-PCR forward primer for *tssK* | Lab stock |
| RT *tssK* RP | CTCACACCCGAACGT ACTGG | qRT-PCR reverse primer for *tssK* | Lab stock |
| RT *tssL* FP | TAACCCAGCAAGAC CCAAGC | qRT-PCR forward primer for *tssL* | This study |
| RT *tssL* RP | TCGCTCTTTTCCACG ACTACG | qRT-PCR reverse primer for *tssL* | This study |
| RT *vgrG* FP | TGACCGTCCGTTTGT AGTGG | qRT-PCR forward primer for *vgrG* (A1S_0550) | This study |
| RT *vgrG* RP | TGACCGCATGGCTAC TTTGT | qRT-PCR reverse primer for *vgrG* (A1S_0550) | This study |
| RT *tetR1* FP | ATGCTGTACTGCCTT TGTCTCT | Forward primer to check for *tetR1* in pAB3 | This study |
| RT *tetR1* RP | CCGTTTCGTGGTCCA CACAT | Reverse primer to check for *tetR1* in pAB3 | This study |
| RT *tetR2* FP | CAACCTCTTGGGCCA GTGTG | Forward primer to check for *tetR2* in pAB3 | This study |
| RT *tetR2* RP | GGTCCACGTGCCACT GATAG | Reverse primer to check for *tetR2* in pAB3 | Lab stock |
| RT AbsR1 FP | GGTTAAGTAAAGAA TTTTAAAG | qRT-PCR forward primer for AbsR1 | Lab stock |
| RT AbsR1 RP | CTCTACCGAAGCAA AAGC | qRT-PCR reverse primer for AbsR1 | Lab stock |
| RT AbsR11 FP | AACGTAGCGGTGTC ACATCA | qRT-PCR forward primer for AbsR11 | Lab stock |
| RT AbsR11 RP | GGTGAAGAGTCCCA TTCCCT | qRT-PCR reverse primer for AbsR11 | Lab stock |
| RT AbsR25 FP | AAATCATGTGTAGG ACCGAG | qRT-PCR forward primer for AbsR25 | Lab stock |

| | | | |
|---|---|---|---|
| RT AbsR25 RP | AAAGCCTACTCAAG AAGCAG | qRT-PCR reverse primer for AbsR25 | Lab stock |
| RT AbsR28 FP | AAGGAGGACATCAT GCCAAC | qRT-PCR forward primer for AbsR28 | Lab stock |
| RT AbsR28 RP | AATTCAAGCATTCGG ACAAG | qRT-PCR reverse primer for AbsR28 | Lab stock |
| RT AbsR29 FP | CGCAGTCAATCAATC AGTGCATTT | qRT-PCR forward primer for AbsR29 | Lab stock |
| RT AbsR29 RP | GATGCAAAGAGCTT GCCAAT | qRT-PCR reverse primer for AbsR29 | Lab stock |
| RT *hfq* FP | CCTTGACTACCACCC TGAGC | qRT-PCR forward primer for *hfq* | Lab stock |
| RT *hfq* RP | TCTACAGTTGTTCCA GCTCGT | qRT-PCR reverse primer for *hfq* | Lab stock |
| RT 16s FP | AGAGGGTGCGAGCG TTAATC | qRT-PCR housekeeping gene forward primer | Lab stock |
| RT 16s RP | GTTAAGCTCGGGGA TTTCAC | qRT-PCR housekeeping gene reverse primer | Lab stock |
| RT A1S_2838 FP | GGCACCATTCGTAG GTGGTT | Forward primer to check the polar effect of ΔAbsR28 upstream | This study |
| RT A1S_2838 RP | AGCGGAATCGTCTTC TTCGG | Reverse primer to check the polar effect of ΔAbsR28 upstream | This study |
| RT A1S_2839 FP | ATCCGGGTCTTGTCC GAATG | Forward primer to check the polar effect of ΔAbsR28 downstream | This study |
| RT A1S_2839 RP | CCTGAATGGAGCAT CACCCA | Reverse primer to check the polar effect of ΔAbsR28 downstream | This study |
| AbsR28 IVT FP | CCGGAATTCTAATAC GACTCACTATAGGG AGATTTTCAACGGCA C | Forward primer for AbsR28 *in vitro* transcription | This study |
| AbsR28 IVT RP | CCCAAGCTTATTGTC CGAATAGGAATAAA AAAACCTAGCG | Reverse primer for AbsR28 *in vitro* transcription | This study |
| AbsR28-M9 IVT FP | CCGGAATTCTAATAC GACTCACTATAGGG AGATTTTCAACGGCA CTTTTTAAAAA**TCCT** GGACATC | Forward primer for AbsR28-M9 *in vitro* transcription | This study |
| *tssM* IVT FP | TAATACGACTCACTA TAGGGGTTTGCACA AACATCTGTTGAACC A | Forward primer for *tssM in vitro* transcription | This study |
| *tssM* IVT RP | CGTACTCTGCTTGGG TATCCTTTT | Reverse primer for *tssM in vitro* transcription | This study |
| *pntB* IVT FP | CCGGAATTCTAATAC GACTCACTATAGGG GGCCTGCCTGTACTT GG | Forward primer for *pntB in vitro* transcription | This study |
| *pntB* IVT RP | CCCAAGCTTCACCAC CAATCATCCAAATA ACCGG | Reverse primer for *pntB in vitro* transcription | This study |
| MumT-pWBAD-FP | TTGGGCTAGCGAATT CGCTATTTCTTGAAT GCTTTTTATCC | Forward primer for *mumT* complementation | This study |

| | | | |
|---|---|---|---|
| MumT-pWBAD-RP | TACCGAGCTCGAATTCTTAAAGCTTGGTCAAATAATTAAAT | Reverse primer for *mumT* complementation | This study |
| AbsR28-M12 FP | CGATTTGGAAAGATGTCAGCTCACGGACAAG | Forward primer for AbsR28 mutation ($A_{76}A_{78}$) | This study |
| AbsR28-M12 RP | GCTGACATCTTTCCAAATCGTTACTGTGTTTTAC | Reverse primer for AbsR28 mutation ($A_{76}A_{78}$) | This study |
| AbsR28-M14 FP | GTCAGCTTACGGACAAGTGTACAAATTTAATTCACTTG | Forward primer for AbsR28 mutation ($U_{89}$) | This study |
| AbsR28-M14 RP | CACTTGTCCGTAAGCTGACATCGTGCCAAATC | Reverse primer for AbsR28 mutation ($U_{89}$) | This study |
| AbsR28-M15 FP | CAGCTCAAGGTAAAGTGTACAAATTTAATTCAC | Forward primer for AbsR28 mutation ($A_{91}U_{94}A_{95}$) | This study |
| AbsR28-M15 RP | GTACACTTTACCTTGAGCTGACATCGTGCC | Reverse primer for AbsR28 mutation ($A_{91}U_{94}A_{95}$) | This study |
| AbsR28-M20 FP | CACGATGTTTGCAAACGGACAAGTGTACAAATTTAA | Forward primer for AbsR28 mutation ($U_{84}$U85) | This study |
| AbsR28-M20 RP | CTTGTCCGTTTGCAAACATCGTGCCAAATCGTTACTG | Reverse primer for AbsR28 mutation ($U_{84}$U85) | This study |
| *tssM* compensatory F1-FP | ACGACGGCCAGTGCCAAGCTTGTTTGCACAAACATCTGTTGAACC | Forward primer for fragment 1 of *tssM* compensatory mutation | This study |
| *tssM* compensatory F1-RP | AGACATCAATGCCACGAACAATGACAAGGC | Reverse primer for fragment 1 of *tssM* compensatory mutation | This study |
| *tssM* compensatory F2-FP | TGTTCGTGGCATTGATGTCTAGGAATACTTCAATT | Forward primer for fragment 2 of *tssM* compensatory mutation | This study |
| *tssM* compensatory F2-RP | TATGACCATGATTACGAATTCCGTACTCTGCTTGGGTATCCT | Reverse primer for fragment 2 of *tssM* compensatory mutation | This study |
